# Boceprevir, GC-376, and calpain inhibitors II, XII inhibit SARS-CoV-2 viral replication by targeting the viral main protease

**DOI:** 10.1101/2020.04.20.051581

**Authors:** Chunlong Ma, Michael D. Sacco, Brett Hurst, Julia A. Townsend, Yanmei Hu, Tommy Szeto, Xiujun Zhang, Bart Tarbet, Michael T. Marty, Yu Chen, Jun Wang

**Affiliations:** Department of Pharmacology and Toxicology, College of Pharmacy, The University of Arizona, Tucson, AZ, 85721, United States; Department of Molecular Medicine, Morsani College of Medicine, University of South Florida, Tampa, FL, 33612, United States; Institute for Antiviral Research, Utah State University, Logan, UT, 84322, USA; Department of Animal, Dairy and Veterinary Sciences, Utah State University, Logan, UT, 84322, USA; Department of Chemistry and Biochemistry, The University of Arizona, Tucson, AZ, 85721, United States

**Keywords:** SARS-CoV-2, COVID-19, main protease, 3CL protease, calpain inhibitors, boceprevir, GC-376

## Abstract

A novel coronavirus SARS-CoV-2, also called novel coronavirus 2019 (nCoV-19), started to circulate among humans around December 2019, and it is now widespread as a global pandemic. The disease caused by SARS-CoV-2 virus is called COVID-19, which is highly contagious and has an overall mortality rate of 6.96% as of May 4, 2020. There is no vaccine or antiviral available for SARS-CoV-2. In this study, we report our discovery of inhibitors targeting the SARS-CoV-2 main protease (M^pro^). Using the FRET-based enzymatic assay, several inhibitors including boceprevir, GC-376, and calpain inhibitors II, and XII were identified to have potent activity with single-digit to submicromolar IC_50_ values in the enzymatic assay. The mechanism of action of the hits was further characterized using enzyme kinetic studies, thermal shift binding assays, and native mass spectrometry. Significantly, four compounds (boceprevir, GC-376, calpain inhibitors II and XII) inhibit SARS-CoV-2 viral replication in cell culture with EC_50_ values ranging from 0.49 to 3.37 μM. Notably, boceprevir, calpain inhibitors II and XII represent novel chemotypes that are distinct from known M^pro^ inhibitors. A complex crystal structure of SARS-CoV-2 M^pro^ with GC-376, determined at 2.15 Å resolution with three monomers per asymmetric unit, revealed two unique binding configurations, shedding light on the molecular interactions and protein conformational flexibility underlying substrate and inhibitor binding by M^pro^. Overall, the compounds identified herein provide promising starting points for the further development of SARS-CoV-2 therapeutics.

## INTRODUCTION

An emerging respiratory disease COVID-19 started to circulate among human in December 2019. Since its first outbreak in China from an unknown origin, it quickly became a global pandemic. As of May 4, 2020, there are 239,740 deaths among 3,442,234 confirmed cases in 215 countries.^1^ The etiological pathogen of COVID-19 is a new coronavirus, the severe acute respiratory syndrome coronavirus 2 (SARS-CoV-2), also called novel coronavirus (nCoV-2019). As the name indicates, SARS-CoV-2 is similar to SARS, the virus that causes severe respiratory symptoms in human and killed 774 people among 8098 infected worldwide in 2003.^2^ SARS-CoV-2 shares ~82% of sequence identity as SARS and to a less extent for Middle East respiratory syndrome (MERS) (~50%).^3, 4^ SARS-CoV-2 is an enveloped, positive-sense, single-stranded RNA virus that belongs to the β-lineage of the coronavirus.^5^ The β-lineage also contains two other important human pathogens, the SARS coronavirus and MERS coronavirus. The mortality rate of SARS-CoV-2 is around 6.96% as of May 4, 2020, which is lower than that of SARS (~10%) and MERS (~34%).^2^ However, current data indicate that SARS-CoV-2 is more contagious and has a larger R0 value than SARS and MERS,^6^ resulting in higher overall death tolls than SARS and MERS. The SARS-CoV-2 virus is currently spreading at an alarming speed in Europe and the United States.

There is currently no antiviral or vaccine for SARS-CoV-2. The SARS-CoV-2 viral genome encodes a number of structural proteins (e.g. capsid spike glycoprotein), non-structural proteins (e.g. 3-chymotrypsin-like protease (3CL or main protease), papain-like protease, helicase, and RNA-dependent RNA polymerase), and accessary proteins. Compounds that target anyone of these viral proteins might be potential antiviral drug candidates.^7, 8^ In this study, we focus on the viral 3CL protease, also called the main protease (M^pro^), and aim to develop potent M^pro^ inhibitors as SAR-CoV-2 antivirals. The M^pro^ plays an essential role in coronavirus replication by digesting the viral polyproteins at more than 11 sites, and it appears like a high profile target for antiviral drug discovery.^9–12^ The M^pro^ has an unique substrate preference for glutamine at the P1 site (Leu-Gln↓(Ser,Ala,Gly)), a feature that is absent in closely related host proteases, suggesting it is feasible to achieve high selectivity by targeting viral M^pro^. As such, we developed the Fluorescence Resonance Energy Transfer (FRET)-based enzymatic assay for the SARS-CoV-2 M^pro^ and applied it to screen a focused library of protease inhibitors. Here we report our findings of several novel hits targeting SARS-CoV-2 M^pro^ and their mechanism of action. The in vitro antiviral activity of the hits was also evaluated in cell culture using infectious SARS-CoV-2 virus. Overall, our study provides a list of drug candidates for SARS-CoV-2 with a confirmed mechanism of action, and the results might help speed up the drug discovery efforts in combating COVID-19. The compounds identified herein represent one of the most potent and selective SARS-CoV-2 M^pro^ inhibitors so far with both enzymatic inhibition and cellular antiviral activity.^9, 11, 12^ The X-ray crystal structure of SARS-CoV-2 M^pro^ with GC-376 showed that the compound can adapt two configurations R and S, offering a molecular explanation of the high-binding affinity of the aldehyde-containing inhibitors. Significantly, the discovery of calpain II and XII inhibitors as potent SARS-CoV-2 antivirals suggest that it might be feasible to design dual inhibitors against the viral M^pro^ and the host calpains, both of which are important for viral replication.

## RESULTS

### Establishing the FRET-based assay for the SARS-CoV-2 main protease (M^pro^)

The M^pro^ gene from SARS-CoV-2 strain BetaCoV/Wuhan/WIV04/2019 was inserted into pET-29a(+) vector and expressed in BL21(DE3) *E. Coli*. with a His-tag in its C-terminus. The M^pro^ protein was purified with Ni-NTA column to high purity (Fig. 1A). To establish the FRET assay condition, we designed a FRET based substrate using the sequence between viral polypeptide NSP4-NSP5 junction from SARS-CoV-2: Dabcyl-KTSAVLQ/SGFRKME(Edans). We then tested the M^pro^ proteolytic activity in buffers with different pH. We found that M^pro^ displays highest activity in pH 6.5 buffer (Fig. 1B), which contains 20 mM HEPES, 120 mM NaCl, 0.4 mM EDTA, and 4 mM DTT and 20% glycerol. As such, all the following proteolytic assay was conducted using this pH 6.5 buffer. Next, we characterized the enzymatic activity of this SARS-CoV-2 M^pro^ by measuring the Km and Vmax values. When 100 nM M^pro^ was mixed with various concentration of FRET substrate (0 to 200 μM), the initial velocity was measured and plotted against substrate concentration. Curve fitting with Michaelis-Menton equation gave the best-fit values of Km and Vmax as 32.8 ± 3.5 μM and 29.4 ± 1.1 RFU/s, respectively (Fig. 1C).

**Figure 1:**
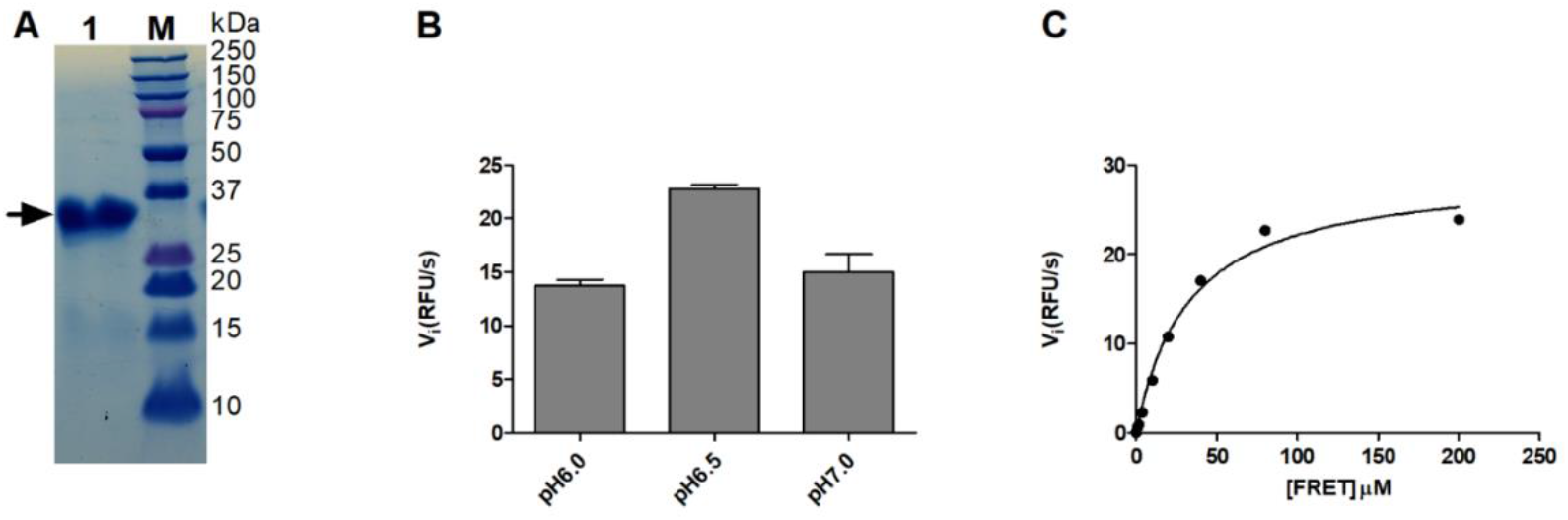
SARS-CoV-2 M^pro^ expression and characterization. (A) SDS-PAGE of His-tagged-Main protease (M^pro^) (lane 1); Lane M, protein ladder; the calculated molecular weight of the His-tagged-M^pro^ is 34,992 Da. (B) Reaction buffer optimization: 250 nM His-tagged-M^pro^ was diluted into three reaction buffers with different pH values. (C) Michaelis-Menten plot of 100 nM His-tagged-M^pro^ with the various concentrations of FRET substrate in pH 6.5 reaction buffer.

**Figure 2:**
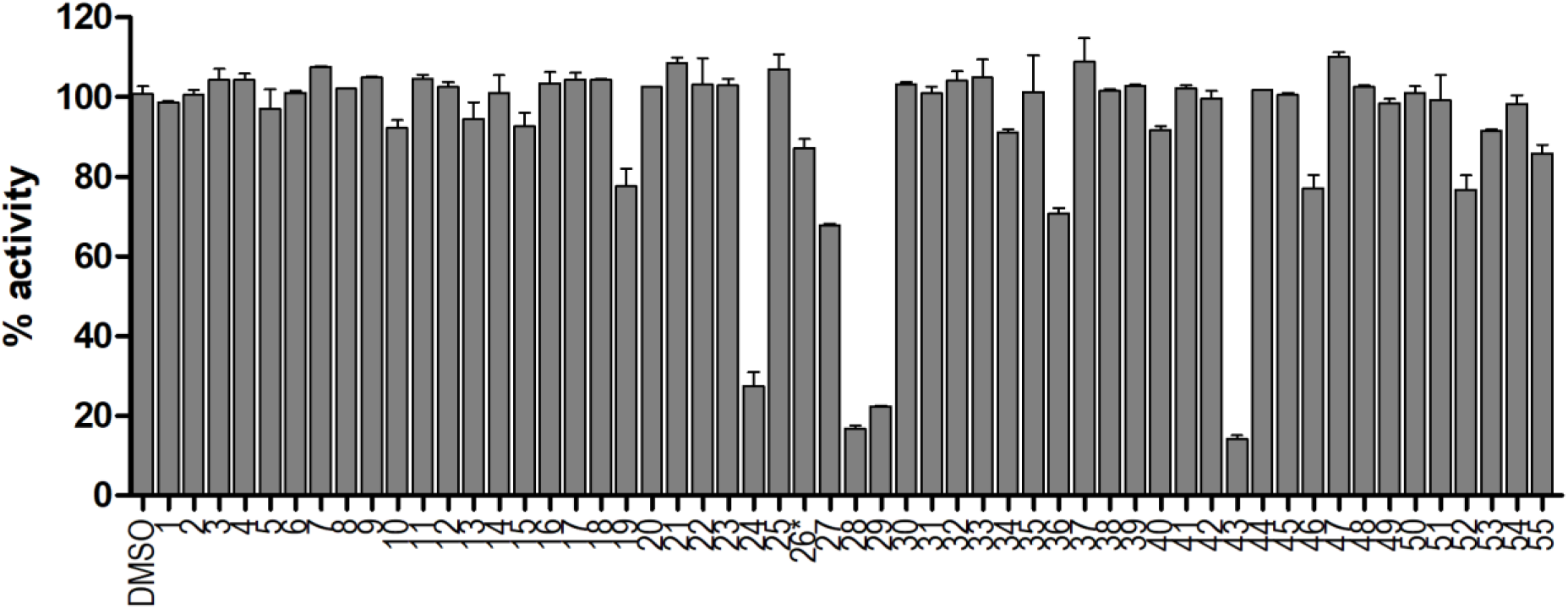
Screening of known protease inhibitors against SARS-CoV-2 M^pro^ using the FRET assay. 20 μM of compounds (**26** was tested at 2 μM) was pre-incubated with 100 nM of SARS-CoV-2 M^pro^ for 30 minutes at 30 °C, then 10 μM FRET substrate was added to reaction mixture to initiate the reaction. The reaction was monitored for 2 hours. The initial velocity was calculated by linear regression using the data points from the first 15 minutes of the reaction. The calculated initial velocity with each compound was normalized to DMSO control. The results are average ± standard deviation of two repeats.

### Primary screening of a focused protease library against the SARS-CoV-2 M^pro^

With the established FRET assay condition, we screened a collection of protease inhibitors from the Selleckchem bioactive compound library to identify potential SARS-CoV-2 M^pro^ inhibitors. The protease inhibitors are grouped based on their targets and mechanism of action and include proteasome inhibitors (**1**-**8**); HIV protease inhibitors (**9**-**14**); γ-secretase inhibitors (**15**-**22**); HCV NS3-4A protease inhibitors (**23**-**29**); DPP-4 inhibitors (**30**-**35**); miscellaneous serine protease inhibitors (**36**-**39**); cathepsin and calpain protease inhibitors (**40**-**43**); miscellaneous cysteine protease inhibitors (**44**-**48**); matrix metalloprotease inhibitors (**49**-**51**); and miscellaneous protease inhibitors (**52**-**55**). The inhibitors were pre-incubated with 100 nM of M^pro^ at 30 °C for 30 minutes in the presence of 4 mM 1,4-dithiothreitol (DTT) before the addition of 10 μM FRET substrate. The addition of DTT was to quench non-specific thiol reactive compounds and to ensure the M^pro^ is in the reducing condition. All compounds were tested at 20 μM, except compound **26**, which was tested at 2 μM due to its fluorescent background. Encouragingly, four inhibitors (**24**, **28**, **29** and **43**) showed more than 60% inhibition against M^pro^ at 20 μM. Among the hits, simeprevir (**24**), boceprevir (**28**), and narlaprevir (**29**) are HCV NS3-4A serine protease inhibitors, and compound MG-132 (**43**) inhibits both proteasome and calpain.

**Table 1.**
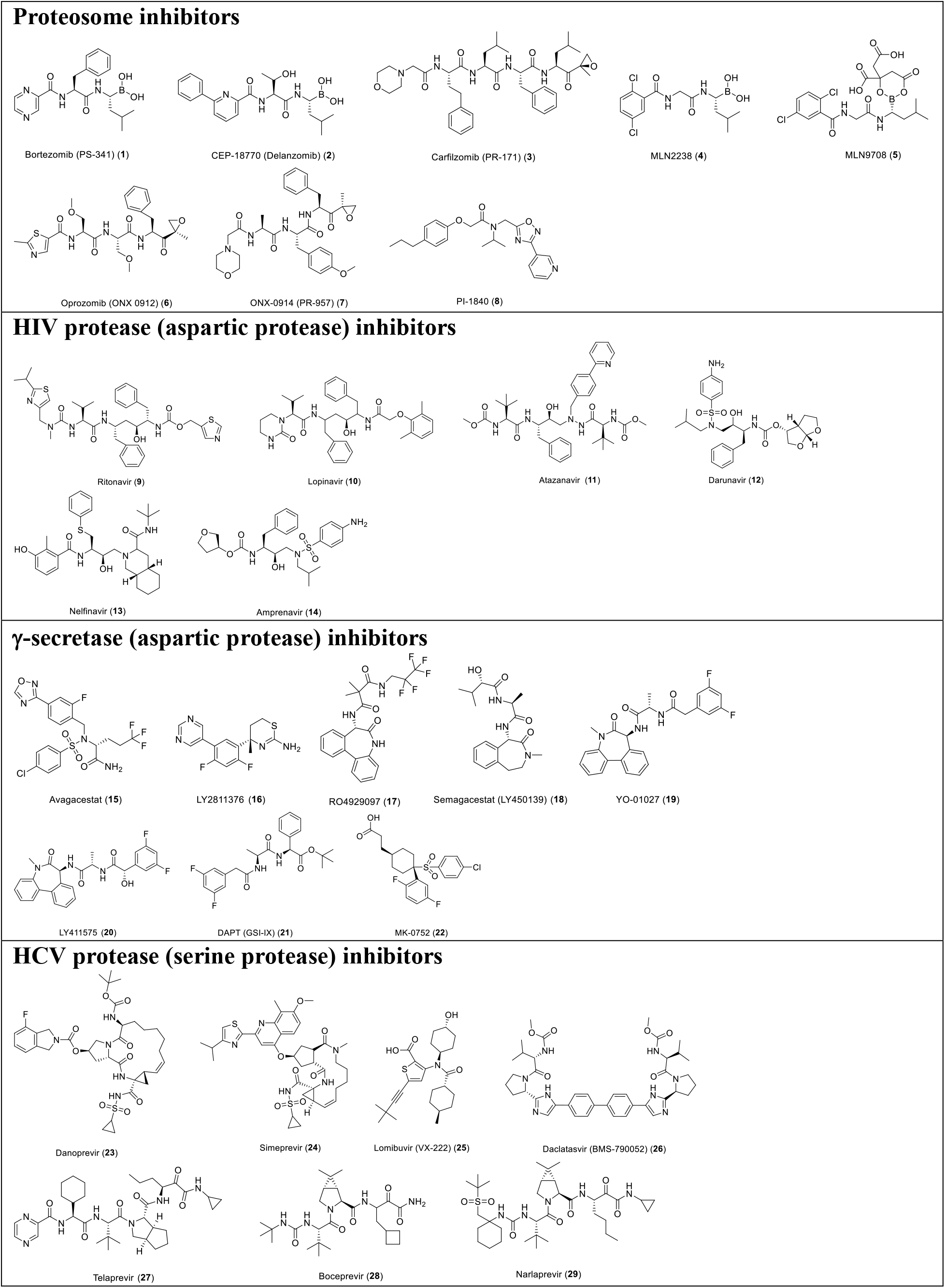
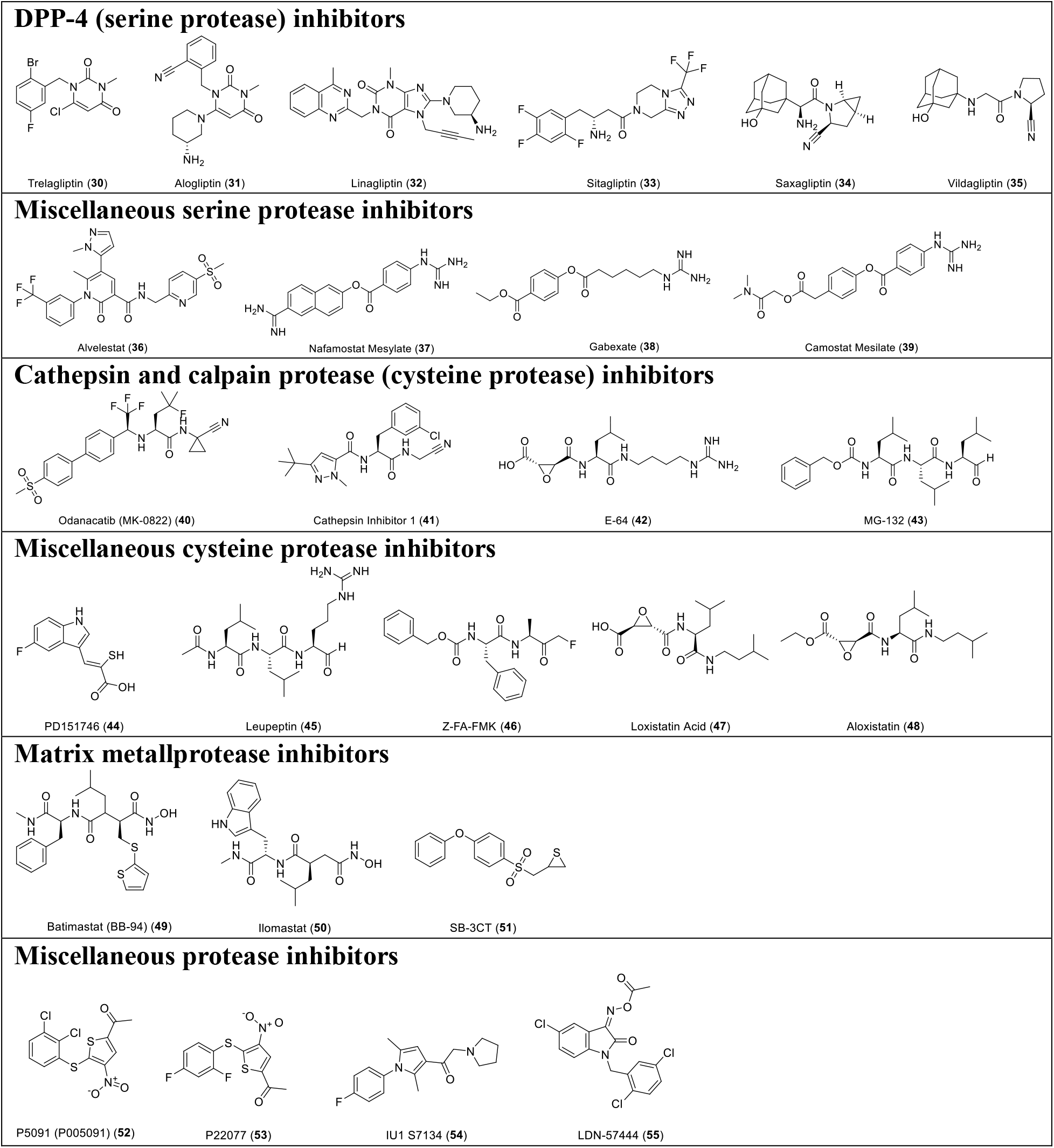
List of protease inhibitors tested against SARS-CoV-2 M^pro^ in the primary FRET assay.

### Secondary screening of a focused library of calpain/cathepsin inhibitors and known viral 3CL^pro^ inhibitors

Given the encouraging results from the primary screening, we then further characterized the four hits (**24**, **28**, **29**, and **43**) in a consortium of assays including dose-response titration, thermal shift binding assay (TSA), and counter screening assays with two other viral cysteine proteases, the enterovirus A71 (EV-A71) 2A and 3C proteases, both of which are cysteine proteases (Table 2). The HCV NS3-4A protease inhibitors boceprevir (**28**) and narlaprevir (**29**) inhibited M^pro^ with IC_50_ values of 4.13 and 4.73 μM, respectively (Table 2), more potent than simeprevir (**24**) (IC_50_ = 13.74 μM). Both compounds **28** and **29** also showed strong binding towards M^pro^ and shifted the melting temperature of the protein (ΔT_m_) by 6.67 and 5.18 °C, respectively, at 40 μM. Despite their potent inhibition against the HCV NS3-4A serine protease and the SARS-CoV-2 cysteine M^pro^, boceprevir (**28**) and narlaprevir (**29**) did not inhibit the EV-A71 2A and 3C proteases (IC_50_ > 20 μM), suggesting they are not non-specific cysteine protease inhibitors. The calpain inhibitor MG-132 (**43**) had an IC_50_ value of 3.90 μM against the M^pro^, and was not active against the EV-A71 2A and 3C proteases (IC_50_ > 20 μM). The binding of MG-132 (**43**) to M^pro^ was also confirmed in the TSA assay with a ΔT_m_ of 4.02 °C.

**Table 2:**
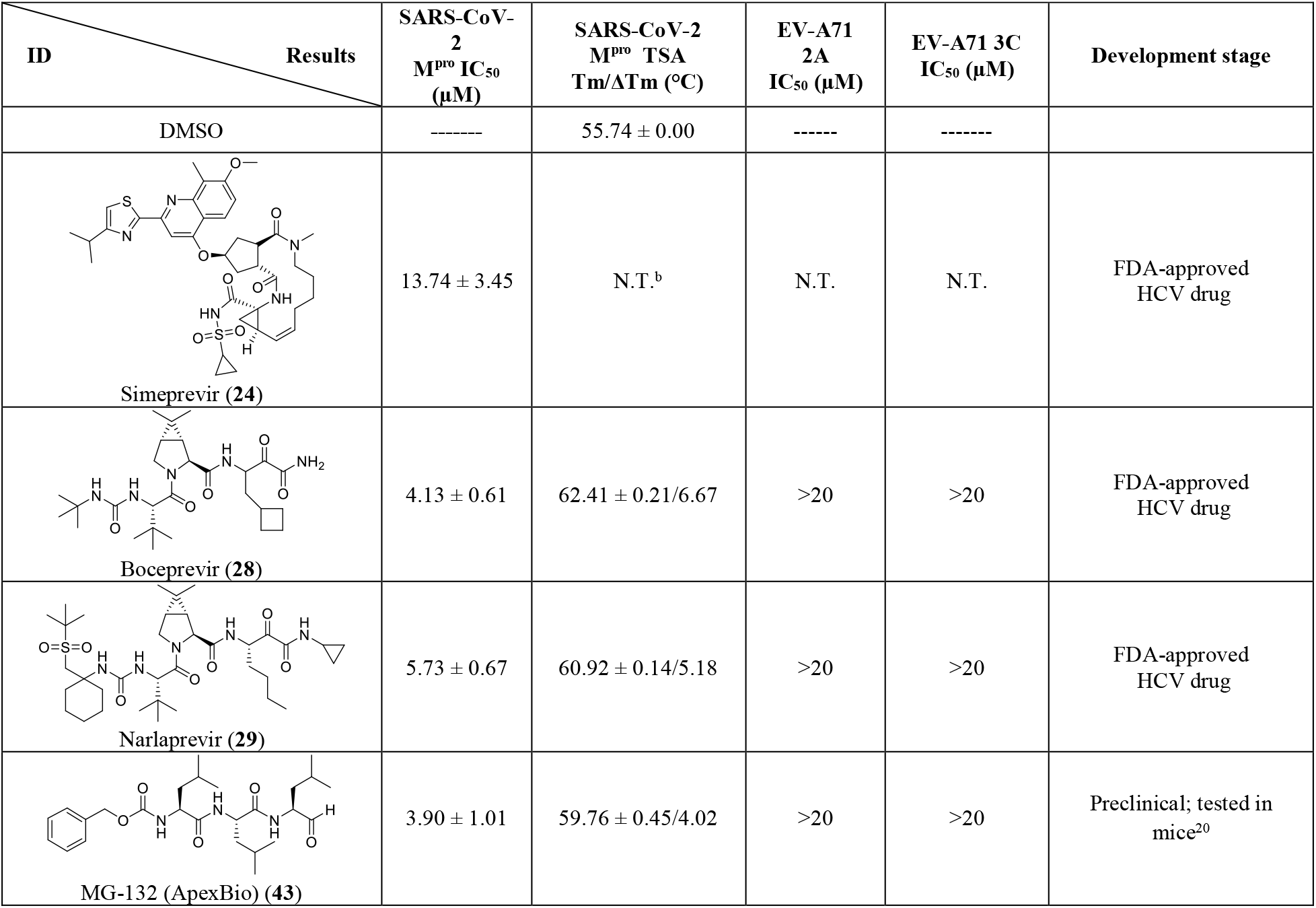
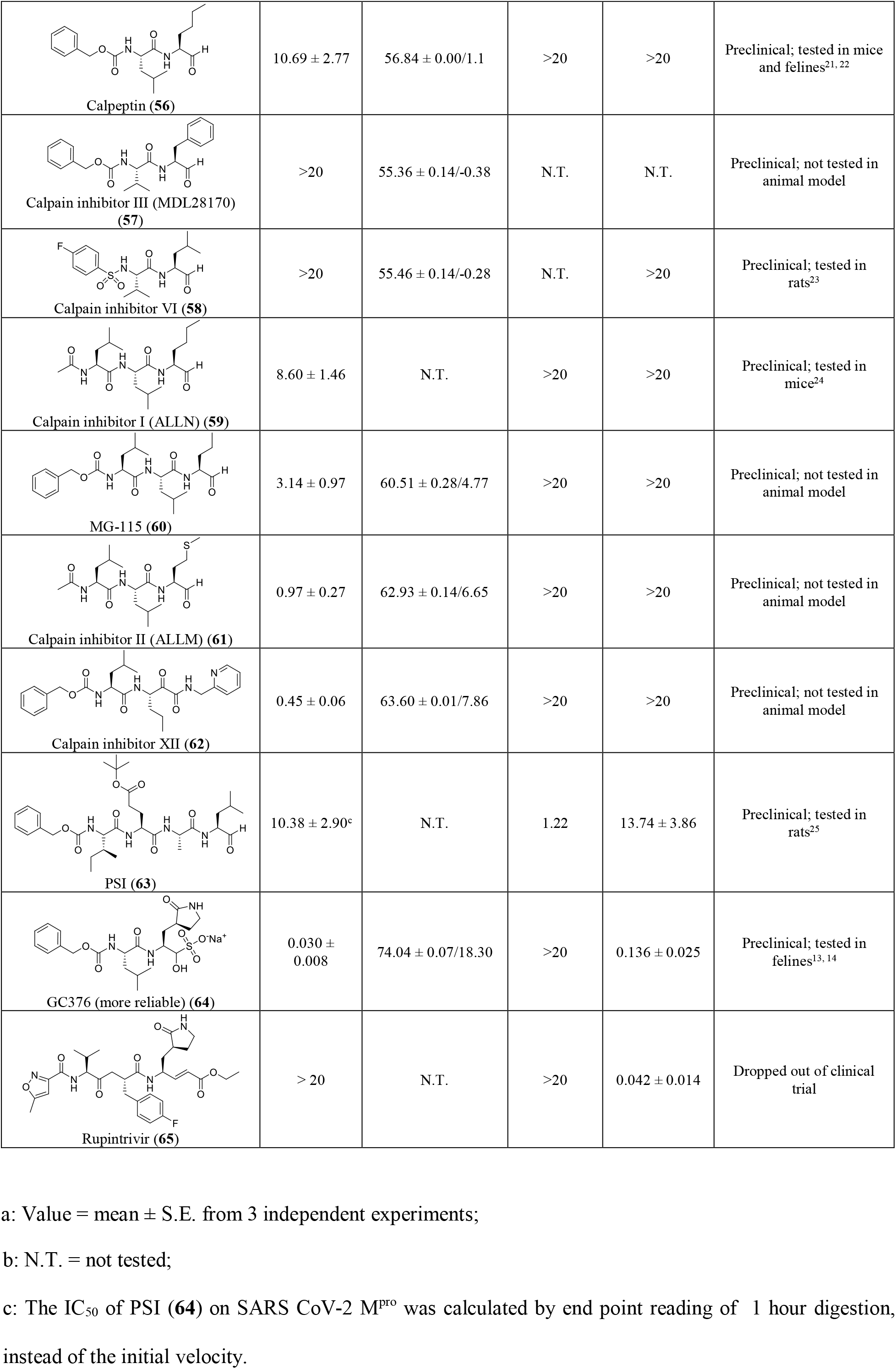
Characterization of HCV and calpain proteases inhibitors against SARS-CoV-2 M^pro^ using a consortium of secondary assays^a^

| ID \ Results | SARS-CoV-2<br>M<sup>pro</sup> IC<sub>50</sub><br>(μM) | SARS-CoV-2<br>M<sup>pro</sup> TSA<br>Tm/ΔTm (°C) | EV-A71<br>2A<br>IC<sub>50</sub> (μM) | EV-A71 3C<br>IC<sub>50</sub> (μM) | Development stage |
| --- | --- | --- | --- | --- | --- |
| DMSO | ----- | 55.74 ± 0.00 | ----- | ----- | |
| Simeprevir (**24**) | 13.74 ± 3.45 | N.T.<sup>b</sup> | N.T. | N.T. | FDA-approved<br>HCV drug |
| Boceprevir (**28**) | 4.13 ± 0.61 | 62.41 ± 0.21/6.67 | >20 | >20 | FDA-approved<br>HCV drug |
| Narlaprevir (**29**) | 5.73 ± 0.67 | 60.92 ± 0.14/5.18 | >20 | >20 | FDA-approved<br>HCV drug |
| MG-132 (ApexBio) (**43**) | 3.90 ± 1.01 | 59.76 ± 0.45/4.02 | >20 | >20 | Preclinical; tested in<br>mice<sup>20</sup> |
| **Calpeptin (56)** | 10.69 ± 2.77 | 56.84 ± 0.00/1.1 | >20 | >20 | Preclinical; tested in mice and felines<sup>21, 22</sup> |
| **Calpain inhibitor III (MDL28170) (57)** | >20 | 55.36 ± 0.14/-0.38 | N.T. | N.T. | Preclinical; not tested in animal model |
| **Calpain inhibitor VI (58)** | >20 | 55.46 ± 0.14/-0.28 | N.T. | >20 | Preclinical; tested in rats<sup>23</sup> |
| **Calpain inhibitor I (ALLN) (59)** | 8.60 ± 1.46 | N.T. | >20 | >20 | Preclinical; tested in mice<sup>24</sup> |
| **MG-115 (60)** | 3.14 ± 0.97 | 60.51 ± 0.28/4.77 | >20 | >20 | Preclinical; not tested in animal model |
| **Calpain inhibitor II (ALLM) (61)** | 0.97 ± 0.27 | 62.93 ± 0.14/6.65 | >20 | >20 | Preclinical; not tested in animal model |
| **Calpain inhibitor XII (62)** | 0.45 ± 0.06 | 63.60 ± 0.01/7.86 | >20 | >20 | Preclinical; not tested in animal model |
| **PSI (63)** | 10.38 ± 2.90<sup>c</sup> | N.T. | 1.22 | 13.74 ± 3.86 | Preclinical; tested in rats<sup>25</sup> |
| **GC376 (more reliable) (64)** | 0.030 ± 0.008 | 74.04 ± 0.07/18.30 | >20 | 0.136 ± 0.025 | Preclinical; tested in felines<sup>13, 14</sup> |
| **Rupintrivir (65)** | > 20 | N.T. | >20 | 0.042 ± 0.014 | Dropped out of clinical trial |

a: Value = mean ± S.E. from 3 independent experiments;
b: N.T. = not tested;
c: The IC<sub>50</sub> of PSI (**64**) on SARS CoV-2 M<sup>pro</sup> was calculated by end point reading of 1 hour digestion, instead of the initial velocity.

In light of the promising results of the calpain inhibitor MG-132 (**43**), we then pursued to testing other calpain and cathepsin inhibitors that are commercially available (**56**-**63**) (Table 2). These compounds were not included in the initial library because they have not been advanced to clinical studies. Among this series of analogs, calpain inhibitor II (**61**) and XII (**62**) are the most potent M^pro^ inhibitors with IC_50_ values of 0.97 and 0.45 μM, respectively. Binding of compounds **61** and **62** to M^pro^ shifted the melting curve of the protein by 6.65 and 7.86 °C, respectively. Encouragingly, both compounds **61** and **62** did not inhibit the EV-A71 2A and 3C proteases (IC_50_ > 20 μM). Calpain inhibitor I (**59**) and MG-115 (**60**) also showed potent inhibition against M^pro^ with IC_50_ values of 8.60 and 3.14 μM, respectively. Calpeptin (**56**) and PSI (**63**) had moderate activity against M^pro^ with IC_50_ values of 10.69 and 10.38 μM, respectively. In contrast, calpain inhibitors III (**57**) and VI (**58**) were not active (IC_50_ > 20 μM).

We also included two well-known viral 3CL protease inhibitors GC-376 (**64**) and rupintrivir (**65**) in the secondary screening. GC-376 (**64**) is an investigational veterinary drug that is being developed for feline infectious peritonitis (FIP).^13, 14^ GC-376 (**64**) was designed to target the viral 3CL protease and had potent antiviral activity against multiple viruses including MERS, FIPV, and norovirus.^13, 15^ Rupintrivir (**65**) was developed as a rhinovirus antiviral by targeting the viral 3CL protease, but it was discontinued in clinical trials due to side effects.^16^ In our study, we found that GC-376 (**64**) was the most potent M^pro^ inhibitor with an IC_50_ value of 0.03 μM. It shifted the melting curve of M^pro^ by 18.30 “C upon binding. In contrast, rupintrivir (**65**) was not active against M^pro^ (IC_50_ > 20 μM). Previous report also showed that rupintrivir was not active against the SARS-CoV 3CL^pr^” (M^pro^) (IC_50_ > 100 μM).^17^ Both compounds **64** and **65** were not active against the EV-A71 2A protease, but showed potent inhibition against the EV-A71 3C protease, which is consistent with previously reported results.^15, 18, 19^

When plotting the IC_50_ values (log scale) of the inhibitors against M^pro^ from the FRET enzymatic assay with the melting temperature shifts (ΔT_m_) from thermal shift binding assay (Fig. 3A), a linear correlation was observed, and the r^2^ of the linear regression fitting is 0.94. This suggests that there is a direct correlation between the enzymatic inhibition and protein binding: a more potent enzyme inhibitor also binds to the protein with higher affinity. The stabilization of the M^pro^ against thermal denaturation was also compound concentration dependent (Fig. 3B).

**Figure 3:**
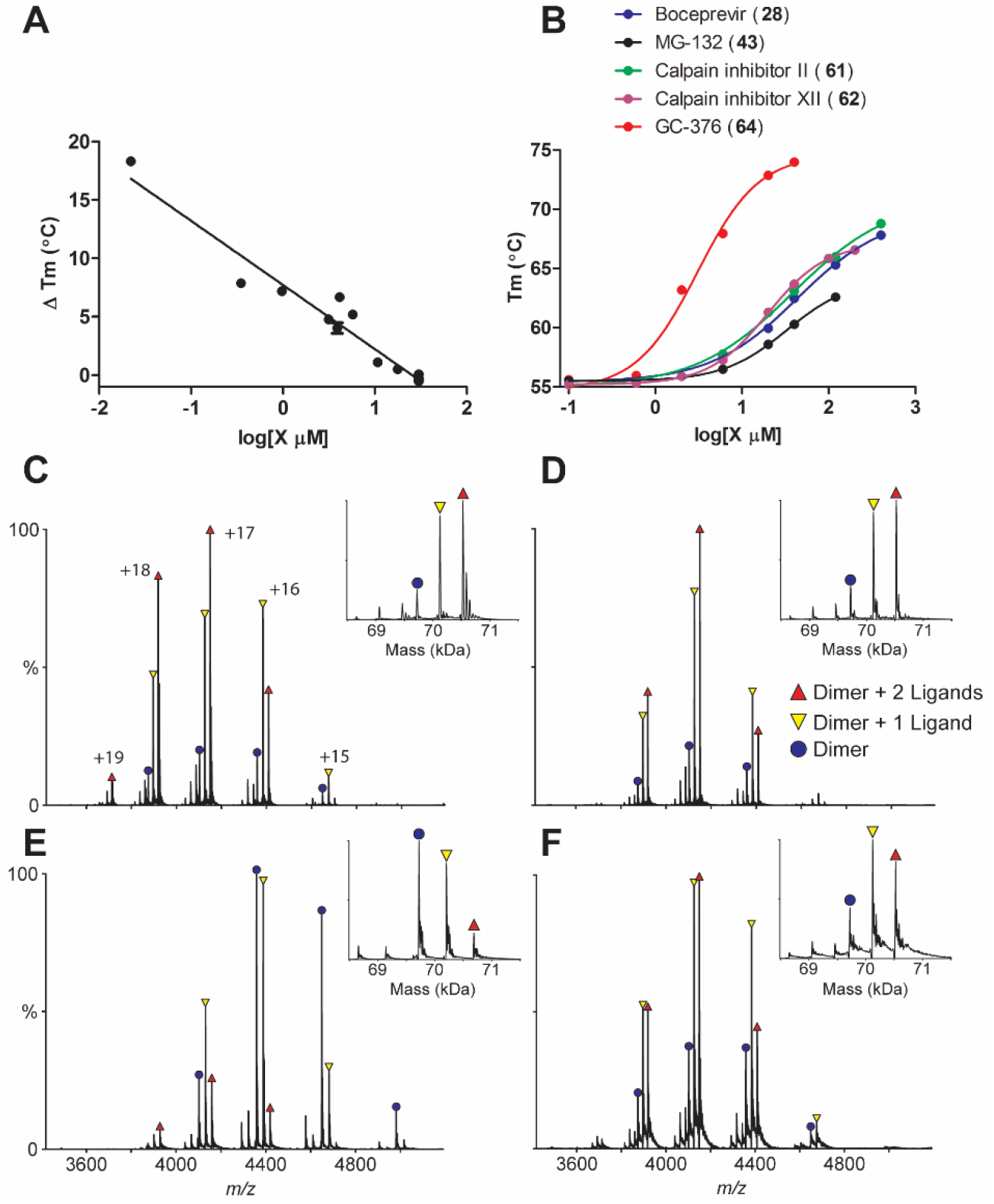
Binding of inhibitors to SARS-CoV-2 M^pro^ using thermal shift binding assay and native mass spectrometry. (A) Correlation of inhibition efficacy (IC_50_) with ΔT_m_ from thermal shift binding assay. Data in Table 2 were used for the plot. The r^2^ of fitting is 0.94. (B) Dose-dependent melting temperature (T_m_) shift. Native MS reveals binding of SARS-CoV-2 M^pro^ to (C) GC-376 (**64**), (D) Calpain inhibitor II (**61**), (E) calpain inhibitor XII (**62**), and (F) Boceprevir (**28**). All ligand concentrations are 12.5 μM except E, which is 25 μM. Peak are annotated for dimer (*blue circle*), dimer with one bound ligand, (*yellow down triangle*), and dimer with two bound ligands (*red up triangle*). Other minor signals are truncated dimers, which bind ligands at the same ratios. Charge states are annotated in C, and insets show the deconvolved zero-charge mass distribution.

The binding of four most potent inhibitors boceprevir (**28**), calpain inhibitors II (**61**), XII (**62**), and GC-376 (**64**) to SARS-CoV-2 M^pro^ was further characterized by native mass spectrometry (MS) (Figs. 3C-F). Native MS analysis showed that the M^pro^ formed a dimer complex with a mass of 69,722 Da, indicating a cleaved N-terminal methionine (Fig. S1). A small amount of monomer and dimer with a single C-terminal truncation of the His-tag was also observed, but the intact dimer was the predominant signal (Fig. S1). Addition of all four ligands tested, boceprevir (**28**), calpain inhibitors II (**61**), XII (**62**), and GC-376 (**64**), showed binding of up to two ligands per dimer (Fig. 3C-F), suggesting a binding stoichiometry of one drug per monomer.

### Mechanism of action of hits

To elucidate the mechanism of action of hits against SARS-CoV-2 M^pro^, we focus on five most potent compounds prioritized from the primary and secondary screenings including boceprevir (**28**), MG-132 (**43**), calpain inhibitor II (**61**), calpain inhibitor XII (**62**), and GC-376 (**64**). For this, we performed enzyme kinetic studies with different concentrations of inhibitors (Fig. 4). A biphasic enzymatic progression curve in the presence but not in the absence of inhibitor is typically a hallmark for a slow covalent binding inhibitor. In the Fig. 4, left column shows the progression curves up to 4 hours. Biphasic progression curves were observed for all 5 inhibitors at high drug concentrations. Significant substrate depletion was observed when the proteolytic reaction proceeded beyond 90 minutes, we therefore chose the first 90 minutes of the progression curves for curve fitting (Fig. 4 middle column). We fit the progression curves in the presence different concentrations of GC-376 (**64**) with the two-step Morrison equation (equation 3 in methods section). GC-376 (**64**) binds to SARS-CoV-2 M^pro^ with an equilibrium dissociation constant for the inhibitor (K_I_) of 59.9 ± 21.7 nM in the first step. After initial binding, a slower covalent bond is formed between GC-376 (**64**) and M^pro^ with the second reaction rate constant (k_2_) being 0.00245 ± 0.00047 s^-1^, resulting an overall k_2_/K_I_ value of 4.08 x 10^4^ M^-1^ s^-1^ (Fig. 4A). However, when we tried to fit the proteolytic progression curves for boceprevir (**28**), MG-132 (**43**), calpain inhibitors II (**61**) and XII (**62**) using the same two-step reaction mechanism, we could not obtain accurate values for the second rate constant k_2_. This is presumably due to significant substrate depletion before the equilibrium between EI and EI*, leading to very small values of k_2_.

**Figure 4:**
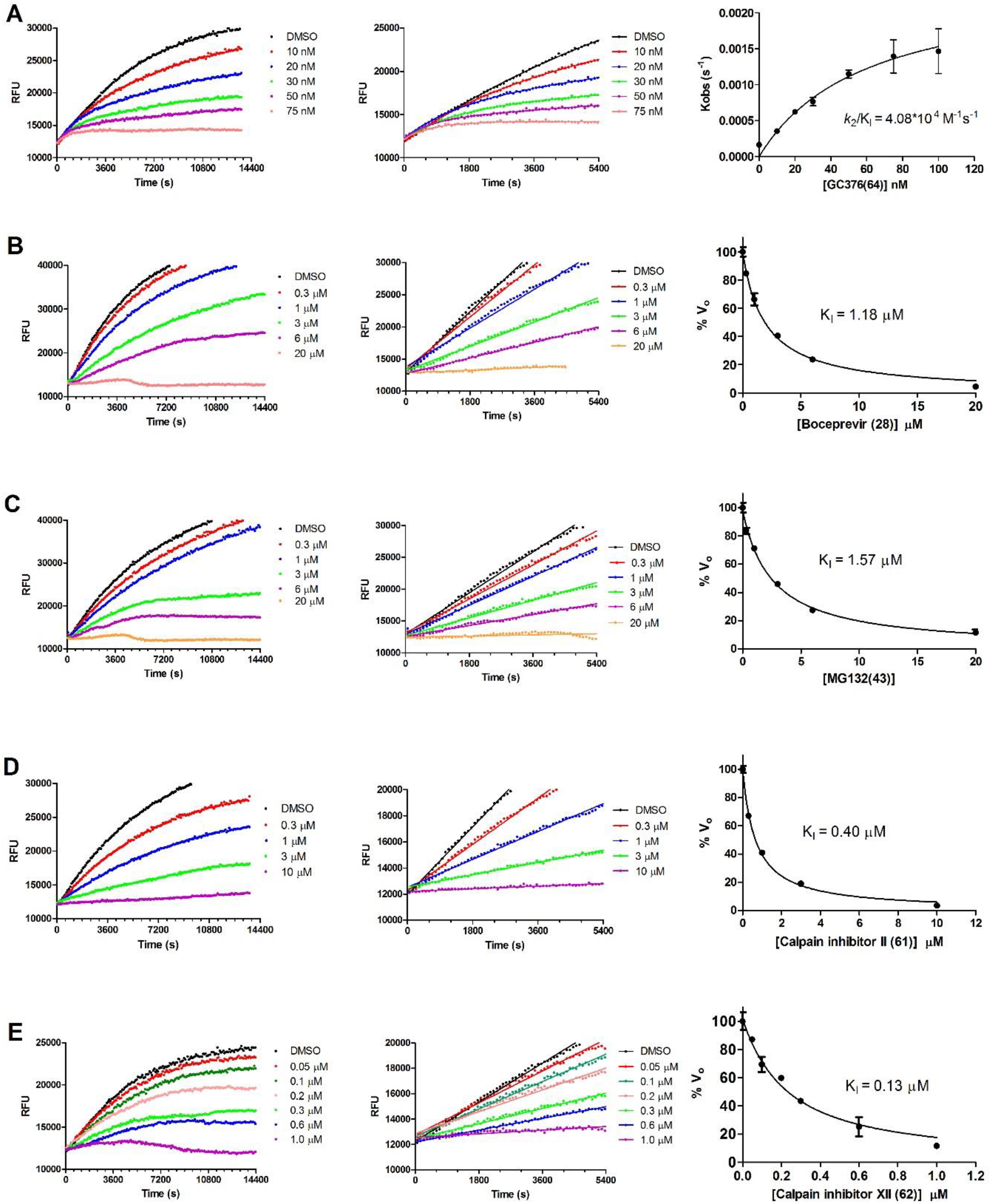
Proteolytic reaction progression curves of M^pro^ in the presence or the absence of compounds. In the kinetic studies, 5 nM M^pro^ was added to a solution containing various concentrations of protease inhibitors and 20 μM FRET substrate to initiate the reaction, the reaction was then monitored for 4 hrs. Left column shows the reaction progression up to 4 hrs; middle column shows the progression curves for the first 90 minutes, which were used for curve fitting to generate the plot shown in the right column. Detailed methods were described in the Method section. (A) GC-376 (**64**); (B) Boceprevir (**28**); (C) MG-132 (**43**); (D) Calpian inhibitor II (**61**); (E) Calpain inhibitor XII (**62**).

Accordingly, for these four inhibitors **28**, **43**, **61**, and **62**, only the dissociation constant K_I_ values from the first step were determined (Figs. 4B-4E). The inhibition constants (K_I_) for boceprevir (**28**), MG-132 (**43**), calpain inhibitors II (**61**) and XII (**62**) are 1.18 ± 0.10 μM, 1.57 ± 0.13 μM, 0.40 ± 0.02 μM, and 0.13 ± 0.02 μM, respectively.

### Cellular antiviral activity and cytotoxicity of hits

To test the hypothesis that inhibiting the enzymatic activity of M^pro^ will lead to the inhibition of SARS-CoV-2 viral replication, we performed cellular antiviral assays for the five promising hits **64**, **28**, **43**, **61**, and **62** against SARS-CoV-2. For this, we first tested the cellular cytotoxicity of these compounds in multiple cell lines (Table S1). GC-376 (**64**), boceprevir (**28**), and calpain inhibitor II (**61**) were well tolerated and had CC50 values of over 100 μM for all the cell lines tested. MG-132 (**43**) was cytotoxic to all the cells with CC50 values less than 1 μM except A549 cells. Calpain inhibitor XII (**62**) had acceptable cellular cytotoxicity with CC50 values above 50 μM for all the cell lines tested (Table S1).

Next, we chose four compounds boceprevir (**28**), calpain inhibitors II (**61**), XII (**62**), and GC-376 (**64**) for the antiviral assay with infectious SARS-CoV-2. MG-132 (**43**) was not included due to its cytotoxicity. Gratifyingly, all four compounds showed potent antiviral activity against SARS-CoV-2 in the primary viral cytopathic effect (CPE) assay with EC_50_ values ranging from 0.49 to 3.37 μM (Table 3). Their antiviral activity was further confirmed in the secondary viral yield reduction (VYR) assay. The most potent compound was calpain inhibitor XII (**62**), which showed EC_50_ of 0.49 μM in the primary CPE assay and EC_90_ of 0.45 μM in the secondary VYR assay. In comparison, remdesivir was reported to inhibit SARS-CoV-2 in the VYR assay with an EC_50_ of 0.77 μM.^26^ None of the compounds inhibited the unrelated influenza virus A/California/07/2009 (H1N1) virus (EC_50_ > 20 μM) (Table S1), suggesting the antiviral activity of the four compounds (boceprevir, calpain inhibitors II, XII, and GC-376) against SARS-CoV-2 is specific. In comparison with recently reported SARS-CoV-2 M^pro^ inhibitors (Table 3), the hits identified herein represent one of the most potent and selective drug candidates with broad chemical diversity.

**Table 3:**
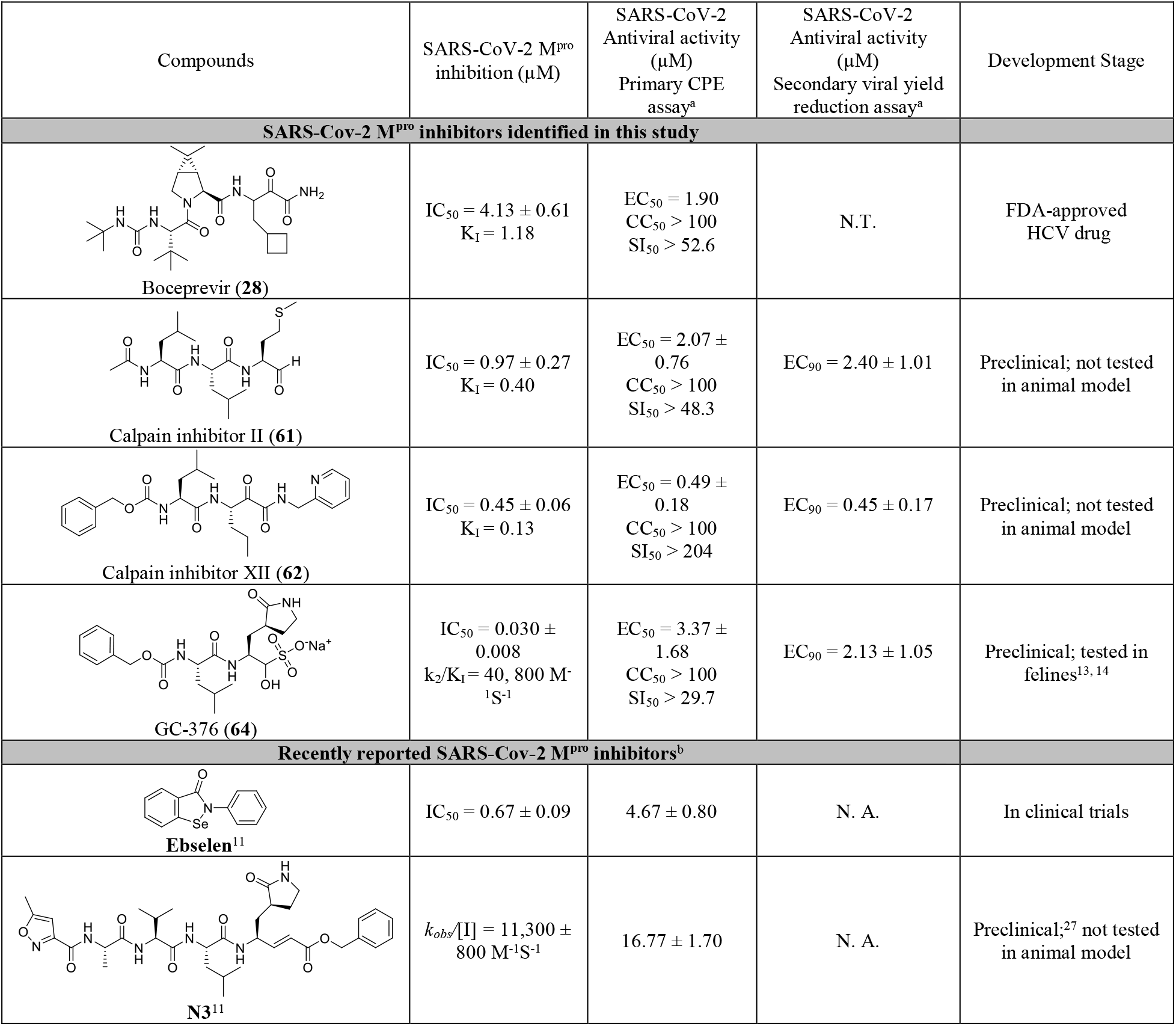
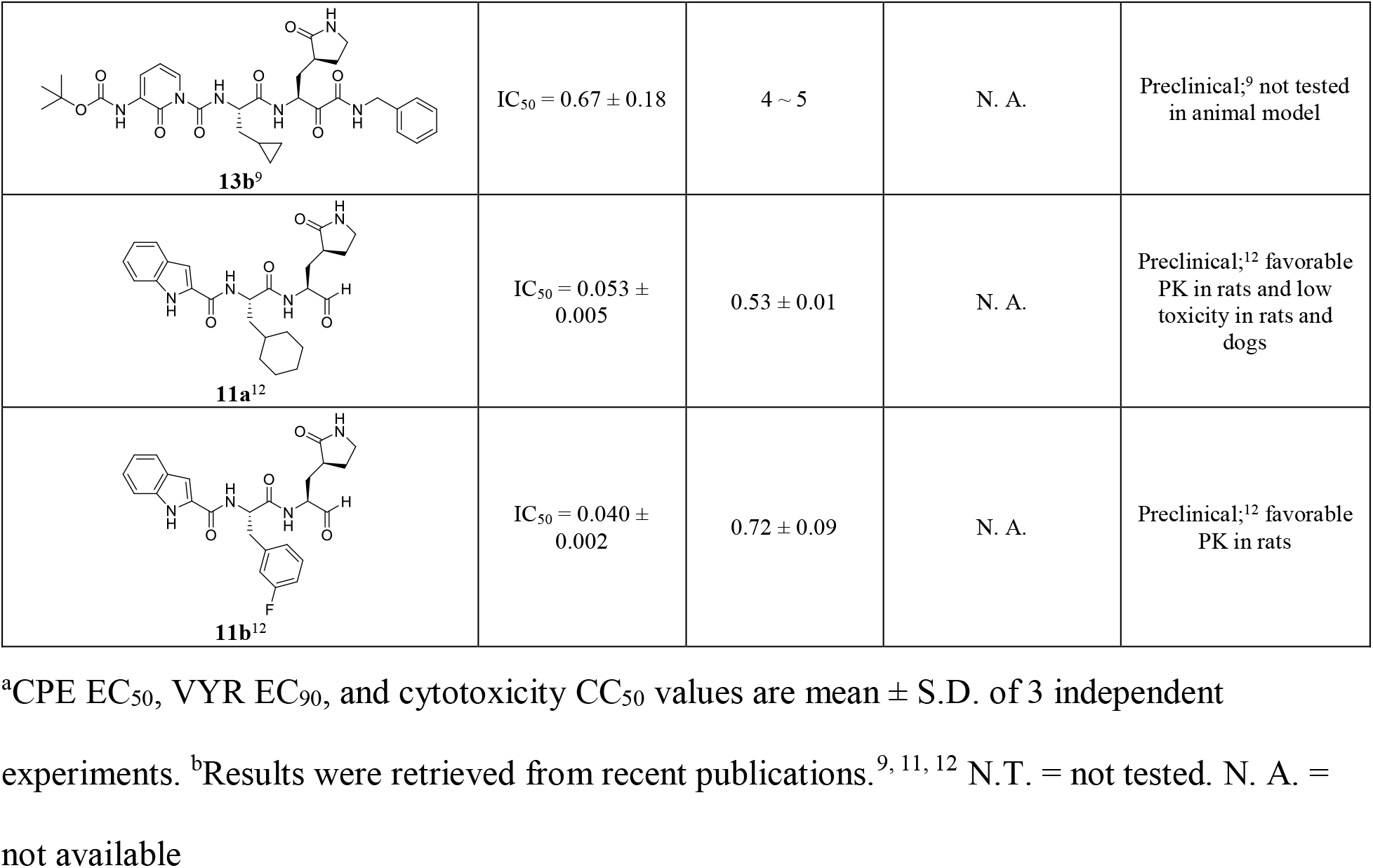
Antiviral activity of hits against SARS-CoV-2 and the comparison with recently reported M^pro^ inhibitors.

| Compounds | SARS-CoV-2 M<sup>pro</sup> inhibition (μM) | SARS-CoV-2 Antiviral activity (μM)<br>Primary CPE assay<sup>a</sup> | SARS-CoV-2 Antiviral activity (μM)<br>Secondary viral yield reduction assay<sup>a</sup> | Development Stage |
| --- | --- | --- | --- | --- |
| **SARS-Cov-2 M<sup>pro</sup> inhibitors identified in this study** |  |  |  |  |
| Boceprevir (**28**) | IC<sub>50</sub> = 4.13 ± 0.61<br>K<sub>i</sub> = 1.18 | EC<sub>50</sub> = 1.90<br>CC<sub>50</sub> > 100<br>SI<sub>50</sub> > 52.6 | N.T. | FDA-approved HCV drug |
| Calpain inhibitor II (**61**) | IC<sub>50</sub> = 0.97 ± 0.27<br>K<sub>i</sub> = 0.40 | EC<sub>50</sub> = 2.07 ± 0.76<br>CC<sub>50</sub> > 100<br>SI<sub>50</sub> > 48.3 | EC<sub>90</sub> = 2.40 ± 1.01 | Preclinical; not tested in animal model |
| Calpain inhibitor XII (**62**) | IC<sub>50</sub> = 0.45 ± 0.06<br>K<sub>i</sub> = 0.13 | EC<sub>50</sub> = 0.49 ± 0.18<br>CC<sub>50</sub> > 100<br>SI<sub>50</sub> > 204 | EC<sub>90</sub> = 0.45 ± 0.17 | Preclinical; not tested in animal model |
| GC-376 (**64**) | IC<sub>50</sub> = 0.030 ± 0.008<br>k<sub>2</sub>/K<sub>i</sub> = 40, 800 M<sup>-1</sup>S<sup>-1</sup> | EC<sub>50</sub> = 3.37 ± 1.68<br>CC<sub>50</sub> > 100<br>SI<sub>50</sub> > 29.7 | EC<sub>90</sub> = 2.13 ± 1.05 | Preclinical; tested in felines<sup>13, 14</sup> |
| **Recently reported SARS-Cov-2 M<sup>pro</sup> inhibitors<sup>b</sup>** |  |  |  |  |
| Ebselen<sup>11</sup> | IC<sub>50</sub> = 0.67 ± 0.09 | 4.67 ± 0.80 | N. A. | In clinical trials |
| N3<sup>11</sup> | k<sub>obs</sub>/[I] = 11,300 ± 800 M<sup>-1</sup>S<sup>-1</sup> | 16.77 ± 1.70 | N. A. | Preclinical;<sup>27</sup> not tested in animal model |
| **13b**<sup>9</sup> | IC<sub>50</sub> = 0.67 ± 0.18 | 4 ~ 5 | N. A. | Preclinical;<sup>9</sup> not tested in animal model |
| **11a**<sup>12</sup> | IC<sub>50</sub> = 0.053 ± 0.005 | 0.53 ± 0.01 | N. A. | Preclinical;<sup>12</sup> favorable PK in rats and low toxicity in rats and dogs |
| **11b**<sup>12</sup> | IC<sub>50</sub> = 0.040 ± 0.002 | 0.72 ± 0.09 | N. A. | Preclinical;<sup>12</sup> favorable PK in rats |

<sup>a</sup>CPE EC<sub>50</sub>, VYR EC<sub>90</sub>, and cytotoxicity CC<sub>50</sub> values are mean ± S.D. of 3 independent experiments. <sup>b</sup>Results were retrieved from recent publications.<sup>9, 11, 12</sup> N.T. = not tested. N. A. = not available

### Complex crystal structure of SARS-CoV-2 M^pro^ with GC-376 (64)

The crystal structure of the SARS-CoV-2 M^pro^ in complex with GC-376 (**64**) was solved in the p3221 space group at 2.15 Å resolution (Table S2). There are three monomers per asymmetric unit (ASU), with two constituting a biological dimer and the third forming a dimer with a crystallographic symmetry related neighboring protomer (Fig. S2). The presence of three monomers in our crystal structure allowed us to capture different binding configurations of GC-376 (**64**) (Fig. 5), a unique feature that was not observed in previous X-ray crystal structures ^9, 11, 12^. The pairwise r.m.s.d among the monomer backbone C α atoms ranges from 0.435 Å to 0.564 Å. Previously, SARS-CoV M^pro^ and SARS-CoV-2 M^pro^ crystal structures have been solved most frequently as a monomer per ASU, and occasionally a dimer ^9, 11, 12, 28, 29^. In its native state, M^pro^ requires dimerization to become catalytically active ^30, 31^, which is supported by our native MS data (Fig. S1). In our crystal structures, all three protomers appear catalytically competent, with the third protomer activated by the N-finger from an adjacent asymmetric unit (Fig. S2B).

**Figure 5:**
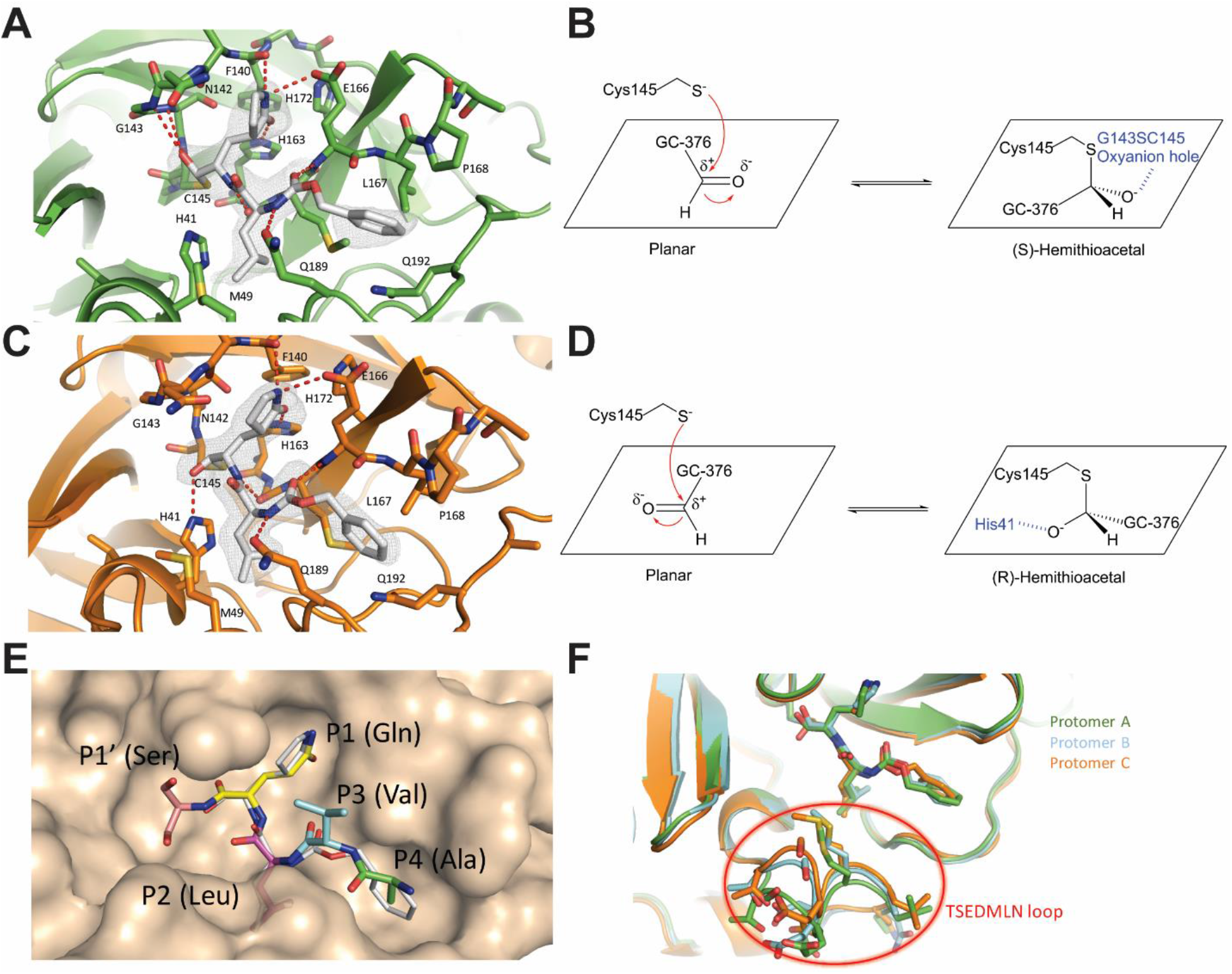
Molecular recognition of GC-376 (64) by SARS-CoV-2 M^pro^. Complex of SARS-CoV-2 M^pro^ and GC-376 (**64**) with (A, B) protomer A and (C, D) protomer C. Unbiased Fo-Fc map, shown in grey, is contoured at 2 σ. Hydrogen bonds are shown as red dashed lines. (E) Surface representation of SARS-CoV-2 M^pro^ + GC-376 (**64**) (white) superimposed with the SARS-CoV M^pro^ natural, N-terminal substrate (PDB ID 2Q6G, with residues P1’-P4 in different colors). The SARS-CoV-2 M^pro^ cleaves between the P1’ and P1 residues. (F) Superimposition of the three protomers in the asymmetric subunit of SARS-CoV-2 M^pro^ with GC-376 (**64**). Significant conformational flexibility is observed, particularly in the TSEDMLN loop.

GC-376 (**64**) forms an extensive network of hydrogen bonds with the active site while also exhibiting excellent geometric complementarity (Fig. 5). These interactions are coupled with the thermodynamic payoff of covalent adduct formation between the aldehyde bisulfite warhead and Cys145, making GC-376 (**64**) one of the most potent SARS-CoV-2 M^pro^ inhibitors *in vitro* with an IC_50_ of 0.030 ± 0.008 μM (Table 3). Along with other known M^pro^ inhibitors **N3**, **13b**, **11a** and **11b** (Table 3), GC-376 (**64**) mimics the peptide substrate that is cleaved by this enzyme (Fig. 5E) ^9, 27, 29, 32^. The glutamine surrogate γ-lactam ring is a cyclized derivative of the P1 glutamine side chain that normally occupies the S1 site; here it forms hydrogen bonds with the His163 and Glu166 side chains and the main chain of Phe140 (Figs. 5A & C). An amide bond connects the γ-lactam side chain to an isobutyl moiety that embeds itself in the hydrophobic S2 site formed by His41, Met49, and Met169. Normally, this S2 site in SARS-CoV-2 M^pro^ can accommodate a variety of hydrophobic substitutions such as isobutyl in GC-376 (**64**) and **N3**, cyclopropyl in **13b**, cyclohexyl in **11a**, and 3-fluorophenyl in **11b** (Table 3) ^33, 34^. A carbamate bond in GC-376 (**64**), which forms hydrogen bonds with the main chain of Glu166 and the side chain of Gln189, connects the P2 isobutyl group to a phenylmethyl ester that interacts with the aliphatic S4 site. Compared with previous inhibitors, the phenylmethyl ester of GC-376 exhibits high complementarity with the S4 site, and the extensive non-polar interactions may contribute significantly to the potency of this compound (Fig. 5E).

Three copies of GC-376 (**64**) were found in the crystal structure, one in each protomer active site (Figs. 5A, 5C, S3). The configurations of G-C376 (**64**) were consistent in protomers A and B, where the thioacetal hydroxide is positioned in the “oxyanion hole” formed by the backbone amides of Gly143, Ser144, and Cys145 (Figs. 5A & S3), resulting in the (S)-configuration. It is noted that aldehydes **11a** and **11b** also bind in the active site of SARS-CoV-2 M^pro^ in the (S)-configuration (PDB: 6M0K and 6LZE) (Figs. S4A).^12^ In protomer C, however, the same hydroxide group orients outwards from the oxyanion hole, forming hydrogen bonds with His41 (Fig. 5C), which gives the (R)-configuration. This (R)-configuration is consistent with the binding mode of α-ketoamide **13b** in the active site of SARS-CoV-2 M^pro^ (PDB: 6Y2F) (Fig. S4B).^9^ These two unique configurations R and S might be a result of the Cys145 thiol nucleophilic attacking the aldehyde from two different faces (Figs. 5B & D). The fact that GC-376 (**64**) can adapt two different configurations R and S upon binding to the active site might explain its high binding affinity towards the target.

An additional difference between the configurations of GC-376 (**64**) in A, B and C is observed in the orientation of the phenylmethyl ester. In protomer C, the CH2 of the phenylmethyl points towards the main chain of Leu167 in a ‘cis’ conformation (Fig. 5C), whereas in protomers A and B this same CH2 points downwards in a ‘trans’ conformation (Figs. 5A & S3). Consequently, this influences the rotameric configuration of the Leu167 isobutyl moiety, where a rotational adjustment of 180° occurs at its “β” carbon. Furthermore, large rearrangements are observed in the flexible TSEDMLN loop consisting of residues 45-51 (TAEDMLN in SARS-CoV-2 M^pro^) that form the S2 and S3’ subsites (Fig. 5F), explaining the broad substrate scope in the P2 site (Table 3). The loop conformations in protomers B and C may be influenced by crystal packing interactions with protomers from adjacent asymmetric units and resemble the conformations in previously determined structures ^11, 12^. Meanwhile, the conformation of protomer A is less restrained and exhibits the most significant conformational divergence. The different loop conformations offer a glimpse of the protein plasticity that allows M^pro^ to accommodate peptides with differing amino acid composition, and underscores the importance of considering this flexibility when analyzing and modeling protein-ligand interactions for M^pro^.

## DISCUSSION

Coronaviruses have caused three epidemics/pandemics in the past twenty years including SARS, MERS, and COVID-19. With the ongoing pandemic of COVID-19, scientists and researchers around the globe are racing to find effective vaccines and antiviral drugs.^7^ The viral polymerase inhibitor remdesivir holds the greatest promise and it is currently being evaluated in several clinical trials.^35, 36^ The HIV drug combination lopinavir and ritonavir recently failed in a clinical trial for COVID-19 with no significant therapeutic efficacy was observed.^37^ To address this unmet medical need, we initiated a drug repurposing screening to identify potent inhibitors against the SARS-CoV-2 M^pro^ from a collection of FDA-approved protease inhibitors. The M^pro^ has been shown to be a validated antiviral drug target for SARS and MERS.^38^ As the SARS-CoV-2 M^pro^ shares a high sequence similarity with SARS and to a less extent with MERS, we reasoned that inhibiting the enzymatic activity of SARS-CoV-2 M^pro^ will similarly prevent viral replication.^9, 11^

Noticeable findings from our study include: 1) Boceprevir (**28**), an FDA-approved HCV drug, inhibits the enzymatic activity of M^pro^ with IC_50_ of 4.13 μM, and has an EC_50_ of 1.90 μM against the SARS-CoV-2 virus in the cellular viral cytopathic effect assay. The therapeutic potential of boceprevir (**28**) should be further evaluated in relevant animal models and human clinic trials. Since boceprevir (**28**) is a FDA-approved drug, the dose, toxicity, formulation, and pharmacokinetic properties are already known, which will greatly speed up the design of follow up studies; 2) GC-376 (**64**), an investigational veterinary drug, showed promising antiviral activity against the SARS-CoV-2 virus (EC_50_ = 3.37 μM). It has the highest enzymatic inhibition against the M^pro^ with an IC_50_ value of 0.03 μM. This compound has promising in vivo efficacy in treating cats infected with FIP, and has favorable in vivo pharmacokinetic properties. Therefore, GC-376 (**64**) is ready to be tested in relevant animal models of SARS-CoV-2 when available. Importantly, the X-ray crystal structure of SARS-CoV-2 M^pro^ in complex with GC-376 (**64**) provides a molecular explanation of the high binding affinity of aldehyde-containing compounds as they can adapt two configurations R and S. The conformational flexibility at the TSEDMLN loop explains the broad substrate scope at the P2 position of M^pro^ inhibitors; 3) Three calpain/cathepsin inhibitors, MG-132 (**43**), calpain inhibitors II (**61**) and XII (**62**), are potent inhibitors of M^pro^ and inhibit SARS-CoV-2 with single-digit to submicromolar efficacy in the enzymatic assay. Calpain inhibitors II (**61**) and XII (**62**) also inhibit SARS-CoV-2 in the CPE assay with EC_50_ values of 2.07 and 0.49 μM, respectively. This result suggests that calpain/cathepsin inhibitors are rich sources of drug candidates for SARS-CoV-2. Indeed, previous studies have shown that calpain and cathepsin are required for the proteolytic processing of the coronavirus S protein, a step that is essential for the viral fusion and genome release during the early stage of viral replication.^39^ Calpain and cathepsin inhibitors such as MDL28170 (calpain inhibitor III)^39^, MG-132^40^, calpain inhibitor VI^41^ have been shown to inhibit SARS-CoV replication in cell culture. Other than the increased potency of targeting both M^pro^ and calpain/cathepsin, an additional benefit of such dual inhibitors might be their high genetic barrier to drug resistance. A significant number of calpain/cathepsin inhibitors have been developed over the years for various diseases including cancer, neurodegeneration disease, kidney diseases, and ischemia/reperfusion injury.^42^ Given our promising results of calpain inhibitors II (**61**) and XII (**62**) in inhibiting the SARS-CoV-2 M^pro^ and their potent antiviral activity in cell culture, it might be worthwhile to repurposing them as antivirals for SARS-CoV-2.

All potent SARS-CoV-2 M^pro^ inhibitors contain reactive warheads such as α-ketoamide (boceprevir (**28**), calpain inhibitor XII (**62**)) or aldehyde (MG-132 (**43**), calpain inhibitor II (**61**)) or aldehyde prodrug, the bisulfite (GC-376 (**64**)). This result suggests that reactive warheads might be essential for SARS-CoV-2 M^pro^ inhibition. The compounds identified in this study represent one of the most potent and selective hits reported so far, and are superior than recently reported SARS-CoV-2 M^pro^ inhibitors **ebselen**, **N3**, and **13b** (Table 3). Calpain inhibitor XII (**62**) had similar potency as the recently disclosed compounds **11a** and **11b** (Table 3).^12^ Notably, calpain inhibitor II (**61**) and XII (**62**) have different chemical scaffolds as GC-376 (**64**), **N3**, **13b**, **11a**, and **11b**, therefore providing new opportunities for designing more potent and selective SARS-CoV-2 M^pro^ inhibitors.

Aside from the above positive results, we also showed that ritonavir (**9**) and lopinavir (**10**) failed to inhibit the SARS-CoV-2 M^pro^ (IC_50_ > 20 μM, Fig. 2), which might explain their lack efficacy in clinical trials for COVID-19.^37^ Camostat (**39**) was recently proposed to inhibit SARS-CoV-2 entry through inhibiting the host TMPRSS2, a host serine protease that is important for viral S protein priming.^43^ However, the antiviral activity of camostat has not been confirmed with infectious SARS-CoV-2 virus. In our study, we found camostat (**39**) has no inhibition against the SARS-CoV-2 M^pro^ (IC_50_ > 20 μM).

In summary, this study identified several potent SARS-CoV-2 M^pro^ inhibitors with potent enzymatic inhibition as well as cellular antiviral activity. Further development based on these hits might lead to clinically useful COVID-19 antivirals. They can be used either alone or in combination with polymerase inhibitors such as remdesivir as a means to achieve potential synergic antiviral effect as well as to suppress drug resistance.

## MATERIALS AND METHODS

Details of materials and methods can be found in the supporting information.

## DATA AVAILABILITY

The structure for SARS-CoV-2 M^pro^ has been deposited in the Protein Data Bank with accession number 6WTT.

## ACKNOWLEDGEMENTS

This research was partially supported by the National Institutes of Health (NIH) (Grant AI147325) and the Arizona Biomedical Research Centre Young Investigator grant (ADHS18-198859) to J. W. B. H. and B. T. thanks for the support from the Respiratory Diseases Branch, National Institute of Allergy and Infectious Diseases, NIH, USA (Contract N01-AI-30048). J. A. T. and M. T. M. were funded by the National Institute of General Medical Sciences and National Institutes of Health (Grant R35 GM128624 to M. T. M.). We thank Michael Kemp for assistance with crystallization and X-ray diffraction data collection. We also thank the staff members of the Advanced Photon Source of Argonne National Laboratory, particularly those at the Structural Biology Center (SBC), with X-ray diffraction data collection. SBC-CAT is operated by UChicago Argonne, LLC, for the U.S. Department of Energy, Office of Biological and Environmental Research under contract DE-AC02-06CH11357.

## AUTHOR CONTRIBUTIONS

J. W. and C. M. conceived and designed the study; C. M. expressed the M^pro^ with the assistance of T. S.; C.M. performed the primary screening, secondary IC_50_ determination, thermal shift-binding assay, and enzymatic kinetic studies; M. S. carried out M^pro^ crystallization and structure determination with the assistance of X. Z, and analyzed the data with Y. C.; B. H. and B. T. performed the SARS-CoV-2 CPE and VYR assay; J. A. T. performed the native mass spectrometry experiments with the guidance from M. T. M.; Y. H. performed the plaque reduction assay with influenza A/California/07/2009 (H1N1) virus; J. W. and Y. C. secured funding and supervised the study; J. W., Y.C., and C. M. wrote the manuscript with the input from others.

## ADDITIONAL INFORMATION

Supplementary information accompanies this paper at

## MATERIALS AND METHODS

### Cell lines and viruses

Human rhabdomyosarcoma (RD); A549, MDCK, Caco-2, and Vero cells were maintained in Dulbecco’s modified Eagle’s medium (DMEM), BEAS2B and HCT-8 cells were maintained in RPMI 1640 medium. Both medium was supplemented with 10% fetal bovine serum (FBS) and 1% penicillin-streptomycin antibiotics. Cells were kept at 37°C in a 5% CO2 atmosphere. The USA_WA1/2020 strain of SARS-CoV-2 obtained from the World Reference Center for Emerging Viruses and Arboviruses (WRCEVA).

### Protein expression and purification

SARS CoV-2 main protease (M^pro^ or 3CL) gene from strain BetaCoV/Wuhan/WIV04/2019 was ordered from GenScript (Piscataway, NJ) in the pET29a(+) vector with E. coli codon optimization. pET29a(+) plasmids with SARS CoV-2 main protease was transformed into competent E. coli BL21(DE3) cells, and a single colony was picked and used to inoculate 10 ml of LB supplemented with 50 g/ml kanamycin at 37°C and 250 rpm. The 10-ml inoculum was added to 1 liter of LB with 50 g/ml kanamycin and grown to an optical density at 600 nm of 0.8, then induced using 1.0 mM IPTG. Induced cultures were incubated at 37 °C for an additional 3 h and then harvested, resuspended in lysis buffer (25 mM Tris [pH 7.5], 750 mM NaCl, 2 mM dithiothreitol [DTT] with 0.5 mg/ml lysozyme, 0.5 mM phenylmethylsulfonyl fluoride [PMSF], 0.02 mg/ml DNase I), and lysed with alternating sonication and French press cycles. The cell debris were removed by centrifugation at 12,000 g for 45 min (20% amplitude, 1 s on/1 s off). The supernatant was incubated with Ni-NTA resin for over 2 h at 4°C on a rotator. The Ni-NTA resin was thoroughly washed with 30 mM imidazole in wash buffer (50 mM Tris [pH 7.0], 150 mM NaCl, 2 mM DTT); and eluted with 100 mM imidazole in 50 mM Tris [pH 7.0], 150 mM NaCl, 2 mM DTT. The imidazole was removed via dialysis or on a 10,000-molecular-weight-cutoff centrifugal concentrator spin column. The purity of the protein was confirmed with SDS-PAGE. The protein concentration was determined via 26OnM absorbance with ε 32890. EV-A71 2Apro and 3Cpro were expressed in the pET28b(+) vector as previously described (*1-3*).

### Peptide synthesis

The SARS-CoV-2 M^pro^ FRET substrate Dabcyl-KTSAVLQ/SGFRKME(Edans) was synthesized by solid-phase synthesis through iterative cycles of coupling and deprotection using the previously optimized procedure.(*4*) Specifically, chemmatrix rink-amide resin was used. Typical coupling condition was 5 equiv of amino acid, 5 equiv of HATU, and 10 equiv of DIEA in DMF for 5 minutes at 80 °C. For deprotection, 5% piperazine plus 0.1 M HOBt were used and the mixture was heated at 80°C for 5 minutes. The peptide was cleaved from the resin using 95% TFA, 2.5% Tris, 2.5% H2O and the crude peptide was precipitated from ether after removal of TFA. The final peptide was purified by preparative HPLC. The purify and identify of the peptide were confirmed by analytical HPLC (> 98% purity) and mass spectrometry. [M+3]^3+^ calculated 694.15, detected 694.90; [M+4]^4+^ calculated 520.86, detected 521.35;

### Native Mass Spectrometry

Prior to analysis, the protein was buffer exchanged into 0.2 M ammonium acetate (pH 6.8) and diluted to 10 μM. DTT was dissolved in water and prepared at a 400 mM stock. Each ligand was dissolved in ethanol and diluted to 10X stock concentrations. The final mixture was prepared by adding 4 μL protein, 0.5 μL DTT stock, and 0.5 μL ligand stock for final concentration of 4 mM DTT and 8 μM protein. Final ligand concentrations were used as annotated. The mixtures were then incubated for 10 minutes at room temperature prior to analysis. Each sample was mixed and analyzed in triplicate.

Native mass spectrometry (MS) was performed using a Q-Exactive HF quadrupole-Orbitrap mass spectrometer with the Ultra-High Mass Range research modifications (Thermo Fisher Scientific). Samples were ionized using nano-electrospray ionization in positive ion mode using 1.0 kV capillary voltage at a 150 °C capillary temperature. The samples were all analyzed with a 1,000–25,000 m/z range, the resolution set to 30,000, and a trapping gas pressure set to 3. Between 10 and 50 V of source fragmentation was applied to all samples to aid in desolvation. Data were deconvolved and analyzed with UniDec.(*5*)

### Enzymatic assays

For reaction condition optimization, 200 μM SARS CoV-2 Main protease was used. pH6.0 buffer contains 20 mM MES pH6.0, 120 mM NaCl, 0.4 mM EDTA, 4 mM DTT and 20% glycerol; pH6.5 buffer contains 20 mM HEPES pH6.5, 120 mM NaCl, 0.4 mM EDTA, 4 mM DTT and 20% glycerol, pH7.0 buffer contains 20 mM HEPES pH7.0, 120 mM NaCl, 0.4 mM EDTA, 4 mM DTT and 20% glycerol. Upon addition of 20 μM FRET substrate, the reaction progress was monitored for 1 hr. The first 15 min of reaction was used to calculate initial velocity (Vi) via linear regression in prism 5. Main protease displays highest proteolytic activity in pH6.5 buffer. All the following enzymatic assays were carried in pH6.5 buffer.

For the measurements of Km/Vmax, screening the protease inhibitor library, as well as IC_50_ measurements, proteolytic reaction with 100 nM Main protease in 100 μl pH6.5 reaction buffer was carried out at 30 °C in a Cytation 5 imaging reader (Thermo Fisher Scientific) with filters for excitation at 360/40 nm and emission at 460/40 nm. Reactions were monitored every 90 s. For Km/Vmax measurements, a FRET substrate concentration ranging from 0 to 200 μM was applied. The initial velocity of the proteolytic activity was calculated by linear regression for the first 15 min of the kinetic progress curves. The initial velocity was plotted against the FRET concentration with the classic Michaelis-Menten equation in Prism 5 software. For the screening protease inhibitor library and IC_50_ measurements, 100 nM Main protease was incubated with protease inhibitor at 30°C for 30 min in reaction buffer, and then the reaction was initiated by adding 10 μM FRET substrate, the reaction was monitored for 1 h, and the initial velocity was calculated for the first 15 min by linear regression. The IC_50_ was calculated by plotting the initial velocity against various concentrations of protease inhibitors by use of a dose-response curve in Prism 5 software. Proteolytic reaction progress curve kinetics measurements with GC376, MG132, Boceprevir, Calpain inhibitor II, and Calpain inhibitor XII used for curve fitting, were carried out as follows: 5 nM Main protease protein was added to 20 μM FRET substrate with various concentrations of testing inhibitor in 200 μl of reaction buffer at 30 °C to initiate the proteolytic reaction. The reaction was monitored for 4 hrs. The progress curves were fit to a slow binding Morrison equation (equation 3) as described previously (*1, 6*):

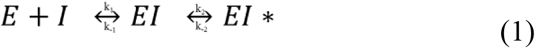

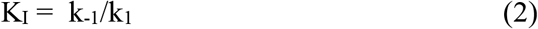

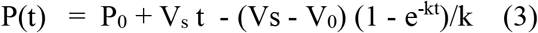

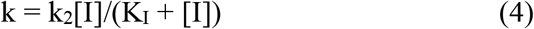

where P(t) is the fluorescence signal at time t, P_0_ is the background signal at time zero, V_0_, V_s_, and and k represent, respectively, the initial velocity, the final steady-state velocity and the apparent first-order rate constant for the establishment of the equilibrium between EI and EI* (*6*). k_2_/K_I_ is commonly used to evaluate the efficacy for covalent inhibitor. We observed substrate depletion when proteolytic reactions progress longer than 90 min, therefore only first 90 min of the progress curves were used in the curve fitting (Figure 6 middle column). In this study, we could not accurately determine the k_2_ for the protease inhibitors: Calpain inhibitor II, MG132, Boceprevir, and Calpain inhibitor XII, due to the very slow k_2_ in these case: significant substrate depletion before the establishment of the equilibrium between EI and EI*. In these cases, K_I_ was determined with Morrison equation in Prism 5.

### Differential scanning fluorimetry (DSF)

The binding of protease inhibitors on Main protease protein was monitored by differential scanning fluorimetry (DSF) using a Thermal Fisher QuantStudio™ 5 Real-Time PCR System. TSA plates were prepared by mixing Main protease protein (final concentration of 3 μM) with inhibitors, and incubated at 30 °C for 30 min. 1× SYPRO orange (Thermal Fisher) were added and the fluorescence of the plates were taken under a temperature gradient ranging from 20 to 90 °C (incremental steps of 0.05 °C/s). The melting temperature (T_m_) was calculated as the mid-log of the transition phase from the native to the denatured protein using a Boltzmann model (Protein Thermal Shift Software v1.3). Thermal shift which was represented as ΔT_m_ was calculated by subtracting reference melting temperature of proteins in DMSO from the Tm in the presence of compound.

### Cytotoxicity measurement

A549, MDCK, HCT-8, Caco-2, Vero, and BEAS2B cells for cytotoxicity CPE assays were seeded and grown overnight at 37 °C in a 5% CO2 atmosphere to ~90% confluence on the next day. Cells were washed with PBS buffer and 200 μl DMEM with 2% FBS and 1% penicillin-streptomycin, and various concentration of protease inhibitors was added to each well. 48 hrs after addition the protease inhibitors, cells were stained with 66 μg/ mL neutral red for 2 h, and neutral red uptake was measured at an absorbance at 540 nm using a Multiskan FC microplate photometer (Thermo Fisher Scientific). The CC50 values were calculated from best-fit dose-response curves using GraphPad Prism 5 software.

### SARS-CoV-2 CPE assay

Antiviral activities of test compounds were determined in nearly confluent cultures of Vero 76 cells. The assays were performed in 96-well Corning microplates. Cells were infected with approximately 60 cell culture infectious doses (CCID50) of SARS-CoV-2 and 50% effective concentrations (EC_50_) were calculated based on virus-induced cytopathic effects (CPE) quantified by neutral red dye uptake after 5 days of incubation. Three microwells at each concentration of compound were infected. Two uninfected microwells served as toxicity controls. Cells were stained for viability for 2 h with neutral red (0.11% final concentration). Excess dye was rinsed from the cells with phosphate-buffered saline (PBS). The absorbed dye was eluted from the cells with 0.1 ml of 50% Sörensen’s citrate buffer (pH 4.2)-50% ethanol. Plates were read for optical density determination at 540 nm. Readings were converted to the percentage of the results for the uninfected control using an Excel spreadsheet developed for this purpose. EC_50_ values were determined by plotting percent CPE versus log10 inhibitor concentration. Toxicity at each concentration was determined in uninfected wells in the same microplates by measuring dye uptake.

### SARS-CoV-2 VYR assay

Virus yield reduction (VYR) assays were conducted by first replicating the viruses in the presence of test compound. Supernatant was harvested 3 days post-infection from each concentration of test compound and the virus yield was determined by endpoint dilution method. Briefly, supernatant virus was serially diluted in log10 increments then plated onto quadruplicate wells of 96-well plates seeded with Vero 76 cells. The presence or absence of CPE for determining a viral endpoint was evaluated by microscopic examination of cells 6 days after infection. From these data, 90% virus inhibitory concentrations (EC_90_) were determined by regression analysis.

### Influenza A virus A/California/07/2009 (H1N1) plaque reduction assay

The plaque assay was performed according to previously published procedures.(*7*)

### M^pro^ crystallization and structure determination

10 mg / mL of SARS-CoV-2 M^pro^ was incubated with 2 mM GC376 at 4° C O/N. The protein was diluted to 2.5 mg / mL the following day in protein buffer (50 mM Tris pH 7.0, 150 mM NaCl, 4 mM DTT). Since GC3760 is water soluble, no precipitation was observed, and centrifugation was not necessary. Crystals were grown by mixing 2 uL of the protein solution with 1 ul of the precipitant solution (15 % PEG 2K, 10% 1,6-hexanediol, and 0.2 M NaCl) in a hanging-drop vapor-diffusion apparatus. Crystals were cryoprotected by transferring to a cryoprotectant solution (20% PEG 2K, 10% 1,6-hexanediol, 20% glycerol) and flash-frozen in liquid nitrogen.

X-ray diffraction data for the SARS-CoV2-M^pro^ GC376 complex structure was collected on the SBC 19-ID beamline at the Advanced Photon Source (APS) in Argonne, IL, and processed with the HKL3000 software suite(*S*). The CCP4 versions of MOLREP were used for molecular replacement using a previously solved SARS-CoV-2 M^pro^ (PDB ID 5RGG) as a reference model(*9*). Rigid and restrained refinements were performed using REFMAC and model building with *COOT(10, 11*). Protein structure figures were made using PyMOL (Schrödinger, LLC).

---

**Supplementary Figure 1.**
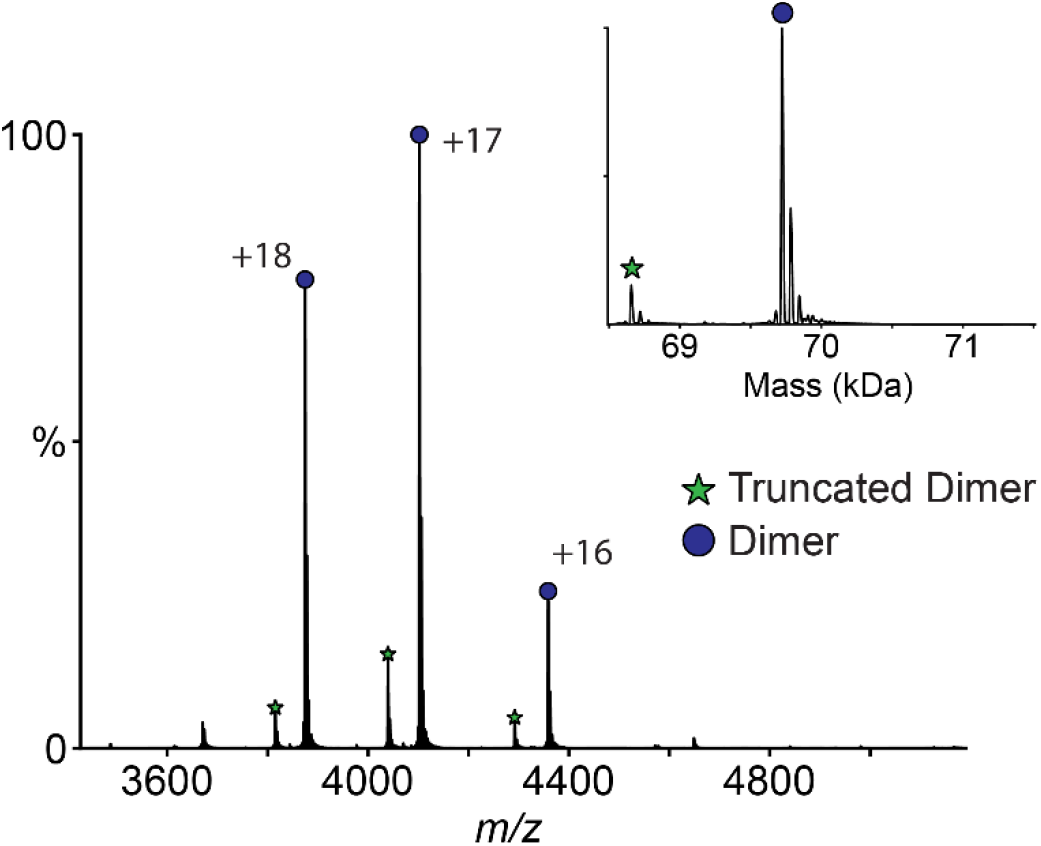
Native mass spectrum of SARS-CoV-2 M^pro^. Native mass spectrum of SARS-CoV-2 VI^1^’^1^’“ with 4 mM DTT shows a dimer (*blue circle*) with a small amount of truncated dimer where one subunit has lost the C-terminal His tag (*green star*). The primary charge states are labeled, and the inset shows the deconvolved zero-charge mass distribution.

**Supplementary Figure 2.**
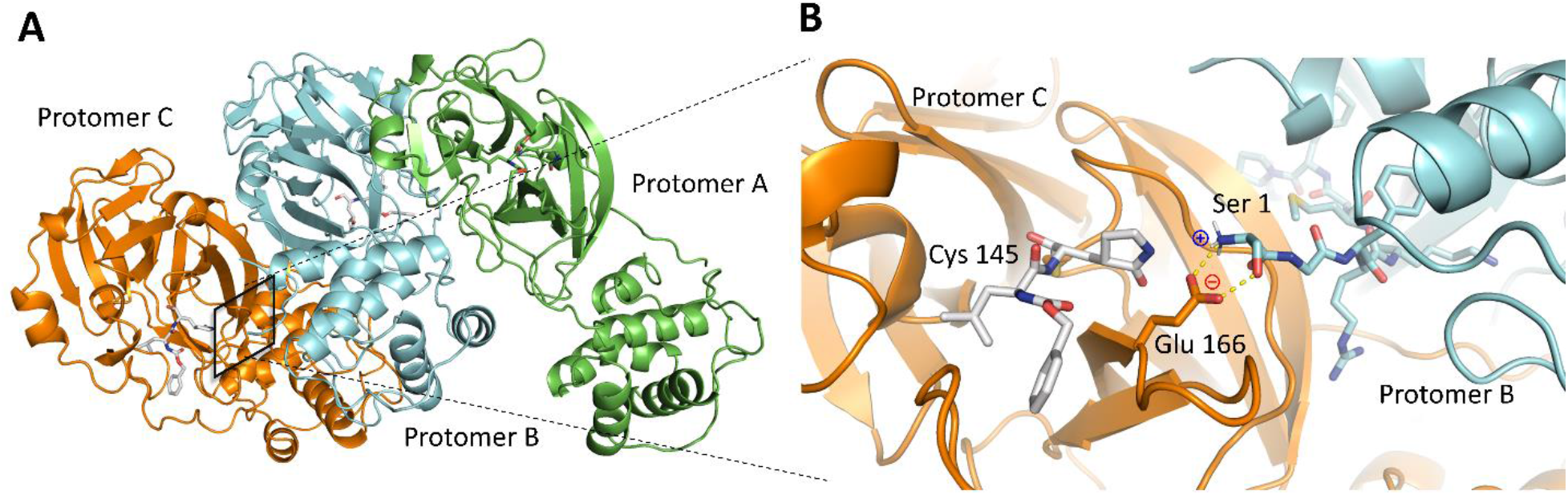
Overall structure of SARS-CoV-2 M^pro^. (A) The three protomers in the asymmetric unit. Protomers B and C form a biological dimer. Protomer A dimerizes with a protomer from an adjacent asymmetric unit (not depicted). (B) The N-finger, or the N-terminal eight residues interact with Glu166 of the adjacent protomer, an important feature for catalytic activity.

**Supplementary Figure 3.**
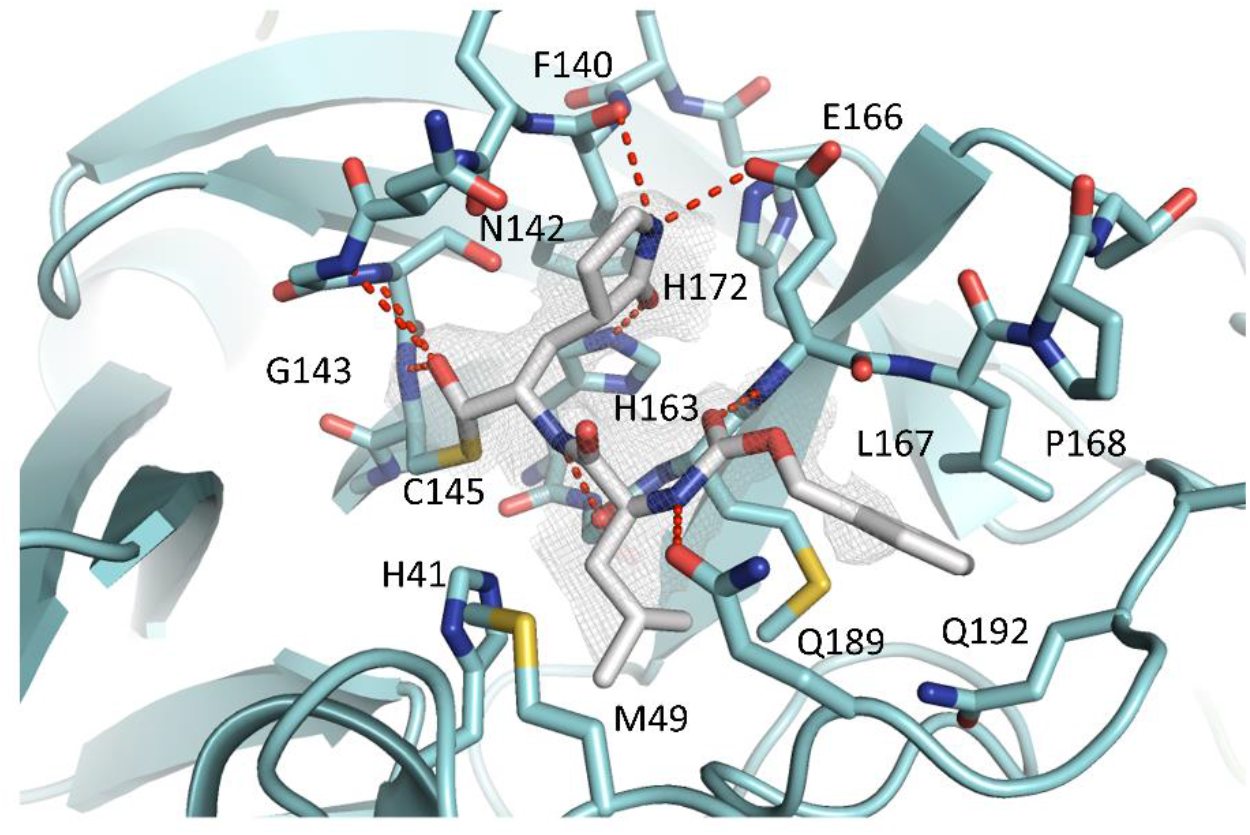
Complex structure of SARS-CoV-2 M^pro^ protomer B with GC-376 (64). Unbiased F_o_-F_c_ map, shown in grey, is contoured at 2 σ. Hydrogen bonds are shown as red dashed lines.

**Supplementary Figure 4.**
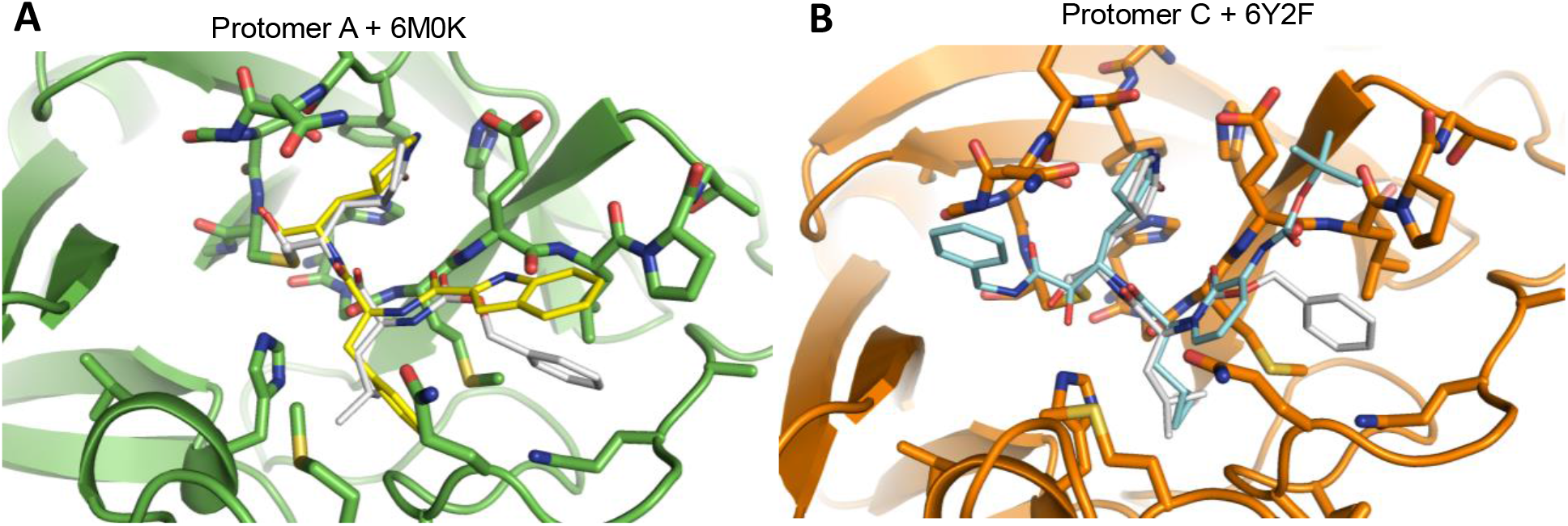
Overlay structures of current X-ray crystal structure with previously solved structures. (A) Overlay structures of protomer A with compound **13b** at the active site (PDB: 6M0K). (B) Overlay structures of protomer C with compound **N3** at the active site (PDB: 6Y2F). Unbiased Fo-Fc map, shown in grey, is contoured at 2 σ. Hydrogen bonds are shown as red dashed lines.

**Supplementary Table 1:** Cytotoxicity of SARS-CoV-2 M^pro^ inhibitors on various cell lines^a^ and the counter screening against influenza virus.

|  | GC-376<br>(64) | Boceprevir<br>(28) | MG-132<br>(43) | Calpain inhibitor II<br>(61) | Calpain inhibitor XII<br>(62) |
| --- | --- | --- | --- | --- | --- |
| MDCK | >100 | >100 | 0.34 ± 0.02 | >100 | 60.36 ± 2.28 |
| Vero | >100 | >100 | 0.45 ± 0.02 | >100 | >100 |
| HCT-8 | >100 | >100 | 0.47 ± 0.02 | >100 | 73.29 ± 11.80 |
| A549 | >100 | >100 | 10.71 ±<br>3.50 | >100 | >100 |
| Caco-2 | >100 | >100 | <0.15 | >100 | 82.02 ± 0.37 |
| BEAS2B | >100 | >100 | 0.14 ± 0.03 | >100 | 78.91 ± 13.70 |
| A/California/07/2009<br>(H1N1) antiviral<br>activity<sup>b</sup> (μM) | > 20 | > 20 | N.T. | > 20 | > 20 |

<sup>a</sup>Cytotoxicity was evaluated by measuring CC<sub>50</sub> values (50% cytotoxic concentration) with CPE assay described in the method section. CC<sub>50</sub> = mean ± S.E. of 3 independent experiments.
<sup>b</sup>Antiviral activity against influenza virus was tested in plaque assay.

**Supplementary Table 2:**
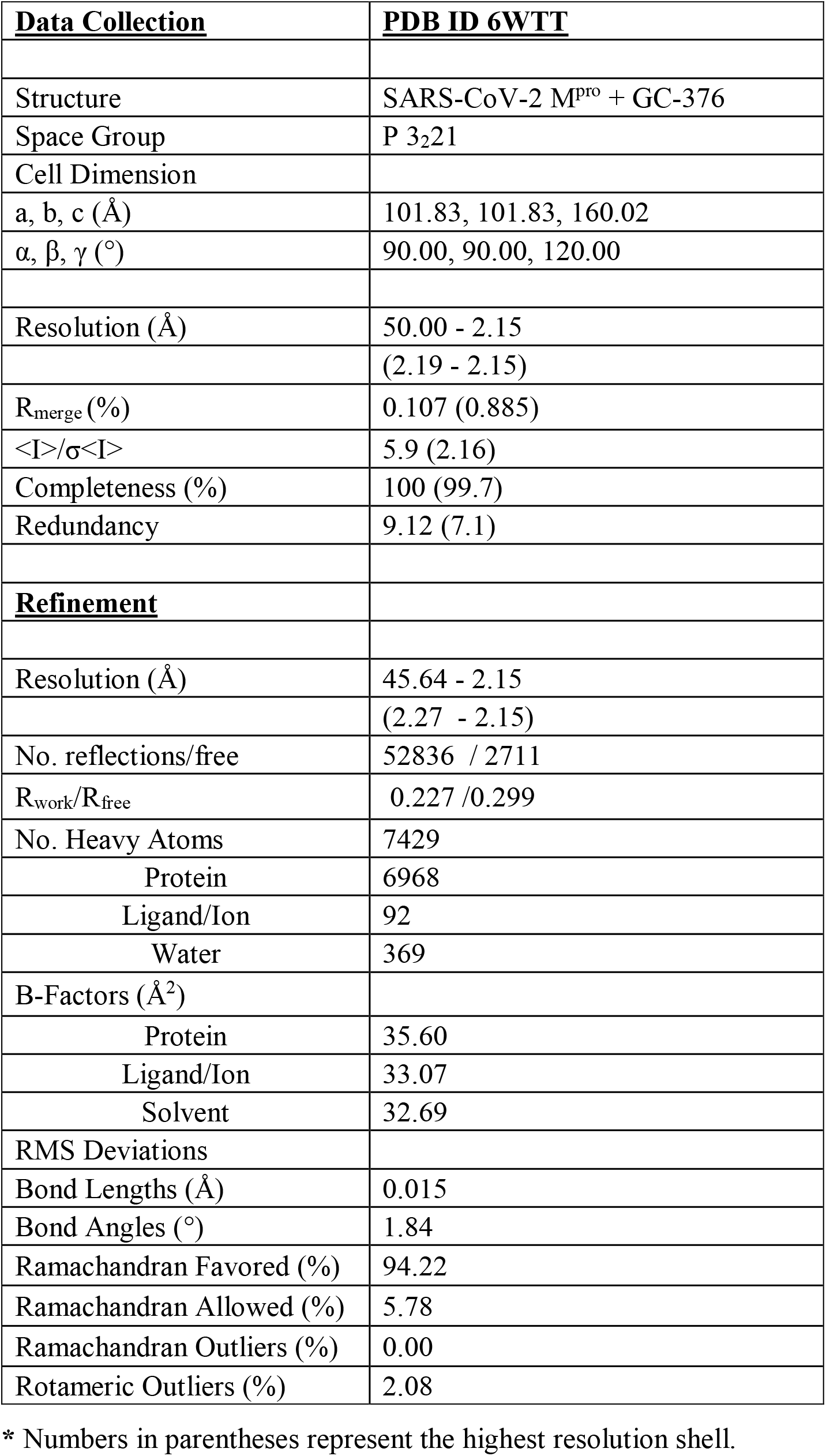
Table of Crystallization Statistics

